# Cross-site reproducibility of human cortical organoids reveals consistent cell type composition and architecture

**DOI:** 10.1101/2023.07.28.550873

**Authors:** Madison R Glass, Elisa A. Waxman, Satoshi Yamashita, Michael Lafferty, Alvaro Beltran, Tala Farah, Niyanta K Patel, Nana Matoba, Sara Ahmed, Mary Srivastava, Emma Drake, Liam T. Davis, Meghana Yeturi, Kexin Sun, Michael I. Love, Kazue Hashimoto-Torii, Deborah L. French, Jason L. Stein

**Author notes:** these authors contributed equally.

## Abstract

**Background:** Reproducibility of human cortical organoid (hCO) phenotypes remains a concern for modeling neurodevelopmental disorders. While guided hCO protocols reproducibly generate cortical cell types in multiple cell lines at one site, variability across sites using a harmonized protocol has not yet been evaluated. We present an hCO cross-site reproducibility study examining multiple phenotypes.

**Methods:** Three independent research groups generated hCOs from one induced pluripotent stem cell (iPSC) line using a harmonized miniaturized spinning bioreactor protocol. scRNA-seq, 3D fluorescent imaging, phase contrast imaging, qPCR, and flow cytometry were used to characterize the 3 month differentiations across sites.

**Results:** In all sites, hCOs were mostly cortical progenitor and neuronal cell types in reproducible proportions with moderate to high fidelity to the *in vivo* brain that were consistently organized in cortical wall-like buds. Cross-site differences were detected in hCO size and morphology. Differential gene expression showed differences in metabolism and cellular stress across sites. Although iPSC culture conditions were consistent and iPSCs remained undifferentiated, primed stem cell marker expression prior to differentiation correlated with cell type proportions in hCOs.

**Conclusions:** We identified hCO phenotypes that are reproducible across sites using a harmonized differentiation protocol. Previously described limitations of hCO models were also reproduced including off-target differentiations, necrotic cores, and cellular stress. Improving our understanding of how stem cell states influence early hCO cell types may increase reliability of hCO differentiations. Cross-site reproducibility of hCO cell type proportions and organization lays the foundation for future collaborative prospective meta-analytic studies modeling neurodevelopmental disorders in hCOs.

## Introduction

Human cortical organoids (hCOs) have been widely adopted to model neurodevelopmental disorders, but concerns that still remain include non-biological variability in organoid structure, cell type composition and gene expression patterns that may confound the interpretations <u>(L. Wang et al. 2023)</u>. Reproducibility has been assessed by individual research groups, demonstrating consistent cell type production across multiple cell lines for guided hCO differentiation protocols <u>(Yoon et al. 2019; Velasco et al. 2019; Rosebrock et al. 2022; Qian et al. 2018; Sloan et al. 2018; Uzquiano et al. 2022)</u>. To date, hCO reproducibility has not been systematically evaluated across multiple independent labs (also called sites) using harmonized protocols. Cross-site differences in hCO differentiation could be caused by many factors including uncontrolled differences in technical variables like handling or stochastic developmental processes. Identifying aspects of hCO differentiations that are reproducible across sites will enable prospective meta-analysis of neurodevelopmental disorder case-control studies, leading to sample sizes that are difficult to achieve in one group alone.

Current evidence suggests limited cross-site reproducibility of *in vitro* neurodevelopmental disorder models. For example, <u>(Griesi-Oliveira et al. 2020)</u> showed only 11.6% to 1.6% of differentially expressed genes were shared across three different studies of idiopathic autism using induced pluripotent stem cell (iPSC) derived neural cultures. Low sample sizes, small effects, multiple causes leading to one disorder, or site-specific technical differences could all decrease the level of consistency. These problems can be effectively addressed by conducting a prospective meta-analysis, in which researchers differentiate hCOs from patient-derived and control iPSC lines at each site using harmonized protocols, measure reproducible phenotypes, and combine data across sites to increase power and decrease inconsistent false positive results <u>(Button et al. 2013)</u>. Prospective meta-analysis would require collaboration, implementation of a harmonized protocol, and reproducibly measuring the same phenotypes across sites.

2D neuronal cultures of an iPSC line with a presenilin mutation and its isogenic control were previously compared across 5 sites using bulk transcriptomics. Of notable concern, site differences were greater than case-control differences <u>(Volpato et al. 2018)</u>. Three dimensional hCO differentiations have been previously compared across labs using pre-existing single cell-RNA-seq (scRNA-seq) datasets. These studies have compared either consistency in developmental trajectories across hCO protocols <u>(Tanaka et al. 2020; He et al. 2023)</u> or fidelity of hCO cell types to the *in vivo* human brain <u>(Bhaduri et al. 2020; Werner and Gillis 2023)</u>, but could not disambiguate differences caused by cell lines, protocols, or technical artifacts from site-specific implementations of the protocol. Additionally, hCO phenotypes other than scRNA-seq have not been compared across multiple labs. Finally, a meta-analysis approach will involve hCO differentiations using harmonized, rather than site-specific, differentiation protocols.

As part of the iPSC working group within the Intellectual and Developmental Disabilities Research Centers (IDDRC) located across the country, we were in the unique position to undertake an hCO cross site reproducibility reproject. Three different IDDRC sites generated hCOs from PGP1, a publicly available iPSC line <u>(Lee et al. 2009)</u>, using a protocol that was pretested across all sites. By using this harmonized protocol, hCO phenotypes were rigorously assessed using multiple techniques including scRNA-seq, phase contrast and 3D fluorescent microscopy, flow cytometry, and qPCR. While there was some variation in visual quality of the hCOs, organoid size, and gene expression patterns, cell type proportions and structure of the hCOs were consistent across all sites.

## Results

### Study design for multimodal test of hCO reproducibility

To test hCO reproducibility, we chose a widely cited guided protocol that demonstrated transcriptomic fidelity to primary fetal cortex and cortical-wall like organization <u>(Qian et al. 2016; Eiraku et al. 2008)</u>. The three IDDRC sites included Children’s Hospital of Philadelphia (CHOP), Children’s National Hospital (CN), and the University of North Carolina at Chapel Hill (UNC).

The same iPSC line (PGP1), bioreactor parts, and matrigel lot were used to standardize differentiations, allowing us to detect differences in hCO phenotypes due to handling differences across sites. Each site completed at least 4 differentiation replicates, defined as a set of organoids seeded and differentiated at the same time from the same passage of PGP1 (Supplemental table 1). We assessed common phenotypes studied in organoids, including cell-type proportions, gene expression, and structure across time using several assays (Figure 1A). In order to minimize site-specific batch effects, samples were collected at each site and shipped to one site that was designated to perform specific assays and corresponding analyses.

**Figure 1.**
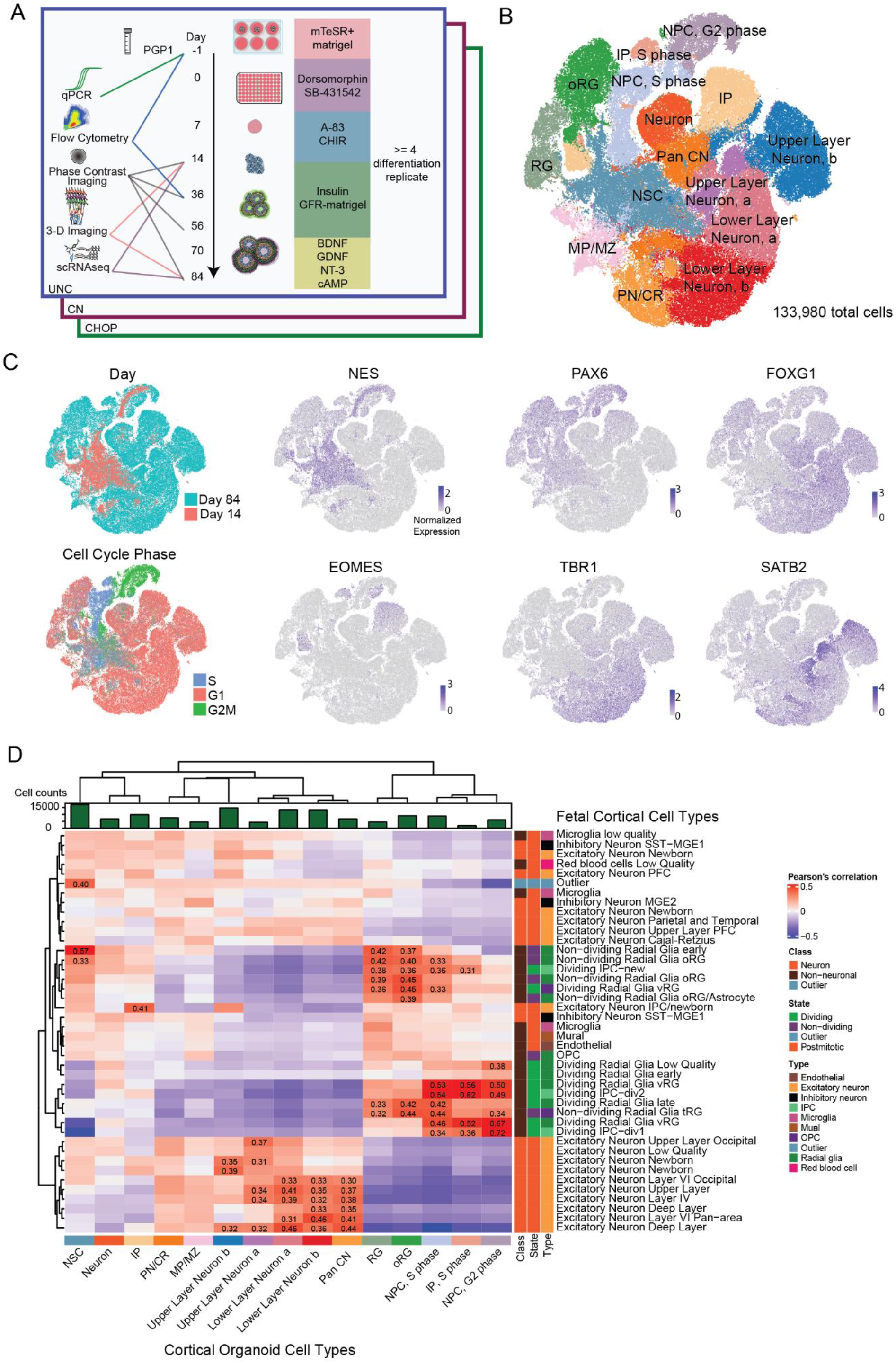
Cell Type Characterizations from Day 14 and Day 84 Human Cortical Organoids. A) Experimental design showing critical steps for differentiation, collection timepoints, and assays. B) tSNE plot of detected cell types across all sites and time points. Abbreviations: NPC, G2: neural progenitor cells, G2-phase. IP, S: intermediate progenitor, S-phase. NPC, S: neural progenitor cell, S-phase. IP: non-dividing intermediate progenitor. oRG: outer radial glia. RG: radial glia. NSC: neuroepithelial stem cell. Neuron: unspecified neuron. Pan CN: pan-cortical neuron. ULN, a: Upper-layer cortical neuron, group a. ULN, b: Upper-layer cortical neuron, group b. LLN, a: Lower-layer cortical neuron, group a. LLN, b: Lower-layer cortical neuron group b. PN/CR: Pan neuron and Cajal–Retzius. MP/MZ: medial pallium and marginal zone. C) tSNE plots colored by differentiation day, inferred cell cycle phase, or gene expression of previously defined markers. D) Organoid cell type correlations to primary fetal telencephalic tissue <u>(Bhaduri et al. 2020)</u>. Pearson’s correlation coefficients above 0.3 and surviving an FDR (Benjamini-Hochberg) adjusted p value threshold of 0.05 are labeled.

### Organoids produce expected cell types

We collected gene expression measurements using a commercially available version of SPLiT-seq <u>(Rosenberg et al. 2018)</u>, yielding 133,980 single cells after quality control (Supplemental Figure 1). hCOs were selected for scRNA-seq based on visible budding at day 14 and smooth edges with expected morphology at day 84 (Supplemental Figure 2). Cells were clustered based on the similarity of their gene expression and then cell-types were annotated using known markers (Figure 1B). hCOs produced expected cell types found in the developing fetal cortex, including neuroepithelial stem cells (NSC), radial glia (RG), intermediate progenitors (IPs), and outer radial glia (oRG), as well as cortical upper and lower layer neurons <u>(Bhaduri et al. 2020; Lui, Hansen, and Kriegstein 2011; Di Lullo and Kriegstein 2017; Kelley and Pașca 2022)</u>.

NSCs expressed *NES* (Lendahl, Zimmerman, and McKay 1990) and *TPM1 (Nicholson-Flynn, Hitchcock-DeGregori, and Levitt 1996)*, with low expression of *PAX6* (Götz, Stoykova, and Gruss 1998) and *FOXG1* (Martynoga et al. 2005), unlike RGs and oRGs (Figure 1C). IPs express the canonical marker *EOMES* (Englund et al. 2005). IP clusters share transcriptomic signatures with neurons (*ROBO1* (Ip et al. 2011), *CUX1 (Coviello et al. 2022)*), as has been previously reported in organoid scRNA-seq studies (Quadrato et al. 2017; Velasco et al. 2019). Some neural progenitor cells (NPCs) expressed S phase markers or G2/M phase markers in addition to *VIM* and *PAX6*. Dividing IPs, delineated by *EOMES* expression and G2/M phase markers, formed a separate cluster from non-dividing IPs.

hCOs produced four cortical neuron populations expressing layer specific markers. Both lower layer neuron clusters expressed *BCL11B* and *TBR1*. Two upper layer neuron clusters were identified: group a expressed *SATB2* and group b expressed *CUX2*. A separate cluster of cortical neurons expressing *DCX* and *FOXG1*, but no layer specific markers, were labeled pan cortical neurons.

We also observed some non-cortical differentiation. hCOs generated a non-cortical cluster with a mix of medial pallium (*TTR* <u>(Pellegrini et al. 2020)</u>) and pioneer neuron markers (*LMX1A* <u>(Yan et al. 2011)</u>). We identified a pan neuron/Cajal-Retzius (PN/CR) cluster expressing gene markers for cortical neurons (*SOX5* <u>(Lai et al. 2008)</u>), ventral brain regions or spinal cord (*PAX3* <u>(Terzić and Saraga-Babić 1999)</u>) and cerebellum (*MEIS1* <u>(Owa et al. 2018)</u>). Interestingly, no expression was detected for markers of the medial ganglionic eminence (*NKX2.1*) and ventral forebrain (*GSX2*), and only 92 cells expressed inhibitory neuron marker *DLX6 - AS* <u>(Bhaduri et al. 2020)</u>. Finally, we labeled an unspecified neuron cluster expressing *SCN2A,* but not *FOXG1* or *VIM*. This cluster shares markers with other hCO “unspecified” neurons <u>(Uzquiano et al. 2022)</u>, such as *GOLGA* gene family members which have poorly defined cellular functions <u>(Bekpen and Tautz 2019)</u>. Non-cortical or poorly defined cells account for only 14% of all cells, demonstrating the majority of hCO cell types were consistent with cortical identity.

To assess the fidelity of hCO cell types to the fetal brain, we correlated cell-type specific hCO gene expression to primary cortical tissue ranging from Carnegie Stage 13 to gestation week 22 <u>(Bhaduri et al. 2020)</u> (Figure 1D). hCO neurons and progenitors generally correlated with their *in vivo* counterparts, whereas cell types absent in the hCOs, like microglia, showed limited correlation. Dividing NPCs showed the highest correlations (r = 0.72), whereas non-dividing cortical progenitors modestly correlated to primary progenitors (r = 0.31 to 0.57).

Interestingly, NSCs correlated to the cell type previously classified as “outlier”, but which likely represents an early progenitor identity because it is only found in Carnegie stage 13 tissue (Figure 2E). The cortical excitatory neuron subtypes identified in hCOs correlated broadly to primary layer-specific cell types (r = 0.3 to 0.46). These findings are similar to <u>(Bhaduri et al. 2020)</u> in that *in vitro* cells represent broad cell classes, but can lack regional or layer specific identities of the *in vivo* brain. The unspecified neuron and PN/CR clusters had correlation coefficients less than 0.3 to all primary cortical cell types, suggesting that these represent non-cortical cell types. Overall, scRNA-seq of hCOs demonstrated that the expected cell types of the developing cortical wall were generated with moderate to high fidelity.

**Figure 2.**
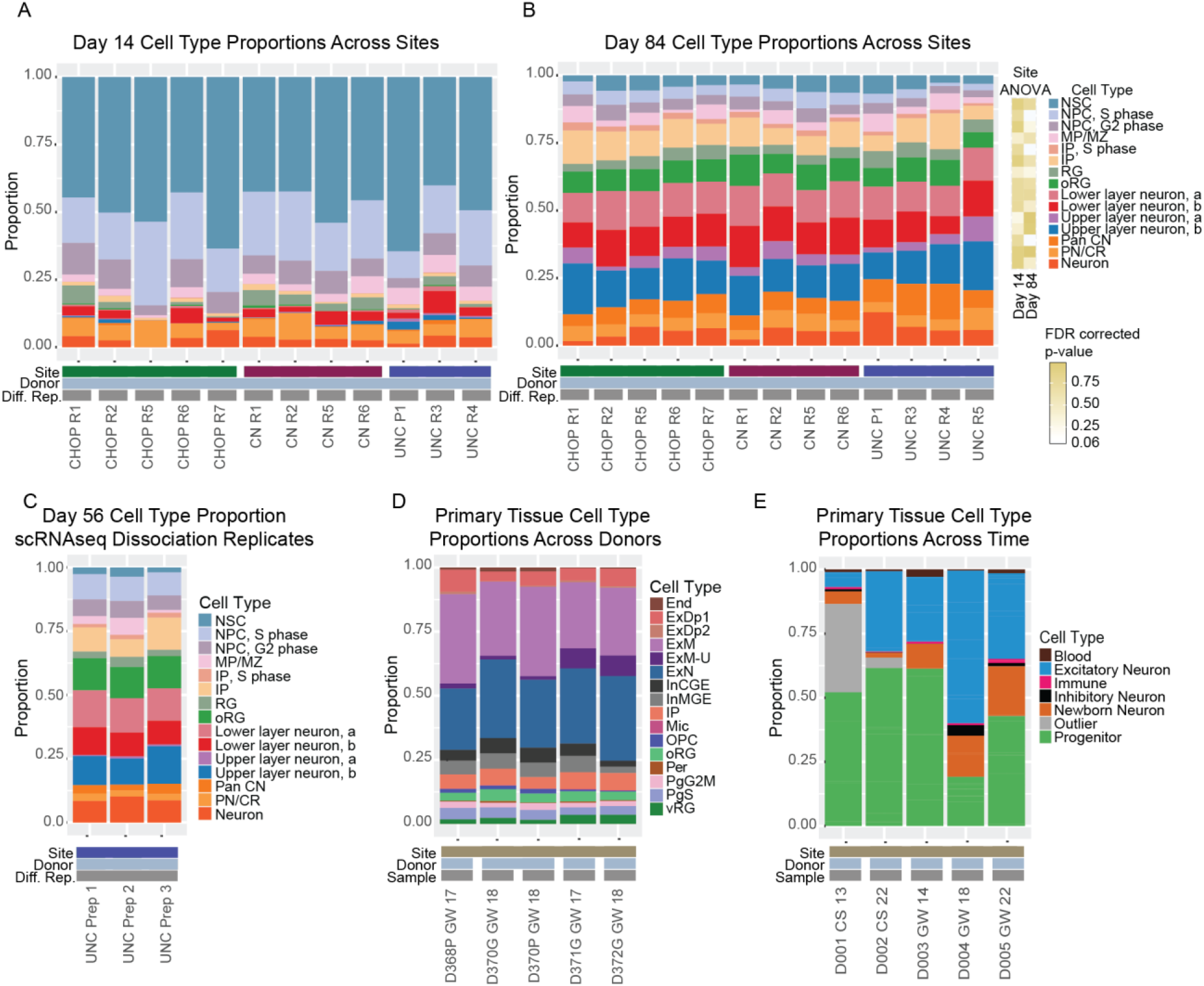
Cell Type Proportions across Sites in Organoid and Primary Tissue Samples. Cell type proportions across the 3 sites at A) day 14 and B) day 84. Each differentiation replicate (Diff. Rep.) is a unique dissociation from a single organoid from an independent differentiation. C) Cell type proportions from 3 separate day 56 organoids harvested from the same differentiation replicate at UNC. D). Cell type proportion reported by <u>(Polioudakis et al. 2019)</u> from primary fetal tissue at similar developmental stages. End (Endothelial), ExDp1 (Excitatory deep layer 1), ExDp2 (Excitatory deep layer 1), ExM (Maturing excitatory), ExM-U (Maturing excitatory upper enriched), ExN (Migrating excitatory), InCGE (Interneuron Caudal Ganglionic Eminence), InMGE (Interneuron Medial Ganglionic Eminence), IP (Intermediate progenitors), Mic (Microglia), OPC (Oligodendrocyte progenitor cell), oRG (Outer radial glia), Per (Pericyte), PgG2M (Cycling progenitors(G2/M phase), PgS (Cycling progenitors (S phase)), vRG (Ventricular radial glia). E) Cell type proportions reported by <u>(Bhaduri et al. 2020)</u>. Each sample is a unique primary tissue donor ordered from earlier to later developmental stages. CS (Carnegie Stage), GW (gestation week).

### Reproducible cell type proportions in hCOs across sites

Next, we calculated cell type proportions at day 14 and day 84 for one hCO from each differentiation replicate to test for cross-site differences using Propeller <u>(Phipson et al. 2022)</u>. Day 14 hCOs were mostly progenitor cell types (>70%), particularly NSCs. No significant differences in cell type proportions across sites were detected at day 14 after correction for multiple comparisons (Supplementary table 2, Figure 2A). However, the PN/CR proportions were nominally different (p-value = 0.018, FDR-corrected p-value = 0.273). At day 84, hCOs are mostly neuronal cell types (>50%). Three cell type proportions were nominally different across sites, but did not survive multiple comparisons correction: RG (p-value = 0.0046, adjusted p-value = 0.061), oRG (p-value = 0.012, adjusted p-value = 0.063), and unspecified neurons (p-value = 0.0081, adjusted p-value = 0.061) (Supplementary Table 2, Figure 2B). Overall, this data suggests high reproducibility of cell type proportions for individual hCOs across differentiation replicates and sites when selecting for visually similar hCOs (Supplementary Figure 2).

In order to quantify cell type variability across hCOs within a single differentiation replicate, three hCOs from the same differentiation replicate at UNC at day 56 were collected (Figure 2C). The coefficient of variation of cell type proportions from multiple organoids in the same differentiation replicate (ranging from 0.778 for medial pallium/marginal zone to 0.0179 for oRG) were comparable to cell type proportions across differentiation replicates at day 84 (ranging from 0.575 for medial pallium/marginal zone to 0.096 for lower layer neuron group a).

Flow cytometry yielded similar measurements of populations of expected telencephalic progenitors (SOX2+/PAX6+, FOXG1+, TBR2+), non-cortical progenitors (GSX2+) and newborn cortical neurons (TBR1+) to the cell type proportions found in the scRNA-seq data (Supplemental Figure 3). No significant cross-site differences were detected, and the cultures had limited neurogenesis within the first month of differentiation.

Finally, we compared hCO cell type proportion variability to primary fetal cortex, which has similar cell type proportions between donors and library preparations. The coefficient of variation ranged from 0.85 for maturing excitatory neurons to 0.12 for excitatory neurons (Figure 2D) <u>(Polioudakis et al. 2019)</u>. Developmental time has the largest effect of cell type proportion across both primary tissue <u>(Bhaduri et al. 2020)</u> and in hCOs (Figure 2A, B, and E). The large differences in cell type composition seen between day 14 and day 84 mimic *in vivo* increases in proportions of neuronal populations and reduced proportions of progenitor populations in the fetal cortex. Together, this data suggests that the variability in cell type proportion of hCOs between differentiation replicates is similar to variability in cell type in primary tissue or library preparation.

### Differential gene expression across sites

To determine transcriptomic differences across sites for each cell type, we performed differential expression on pseudobulked scRNA-seq data at day 14 and day 84 <u>(Love, Huber, and Anders 2014; Crowell et al. 2020)</u>. We detected 786 unique genes with significant differential expression across sites at day 14 (Figure 3A, FDR < 0.05). NSCs, the most abundant cell type, showed the most differentially expressed genes. Differentially expressed genes (DEGs) within abundant cell types like NSC and NPC were enriched in gene ontology (GO) terms implicating tissue development, suggesting small but detectable differences in early differentiation processes across sites (Figure 3C). Specific genes in the NSC simply different levels of regionalization and WNT activity across sites (Figure 3E). For example, *LHX2* had lower expression at UNC. *LHX2* specifies cortical identity in early development <u>(Mangale et al. 2008)</u>. UNC also had higher expression of *RSPO2*, which can bind to LGR4/5 to stabilize WNT signaling <u>(Carmon et al. 2011)</u>. Slight changes in WNT activation or sensitivity across sites may have led to the enrichment of these neurodevelopmental GO terms within DEGs (Harrison-Uy and Pleasure 2012).

**Figure 3.**
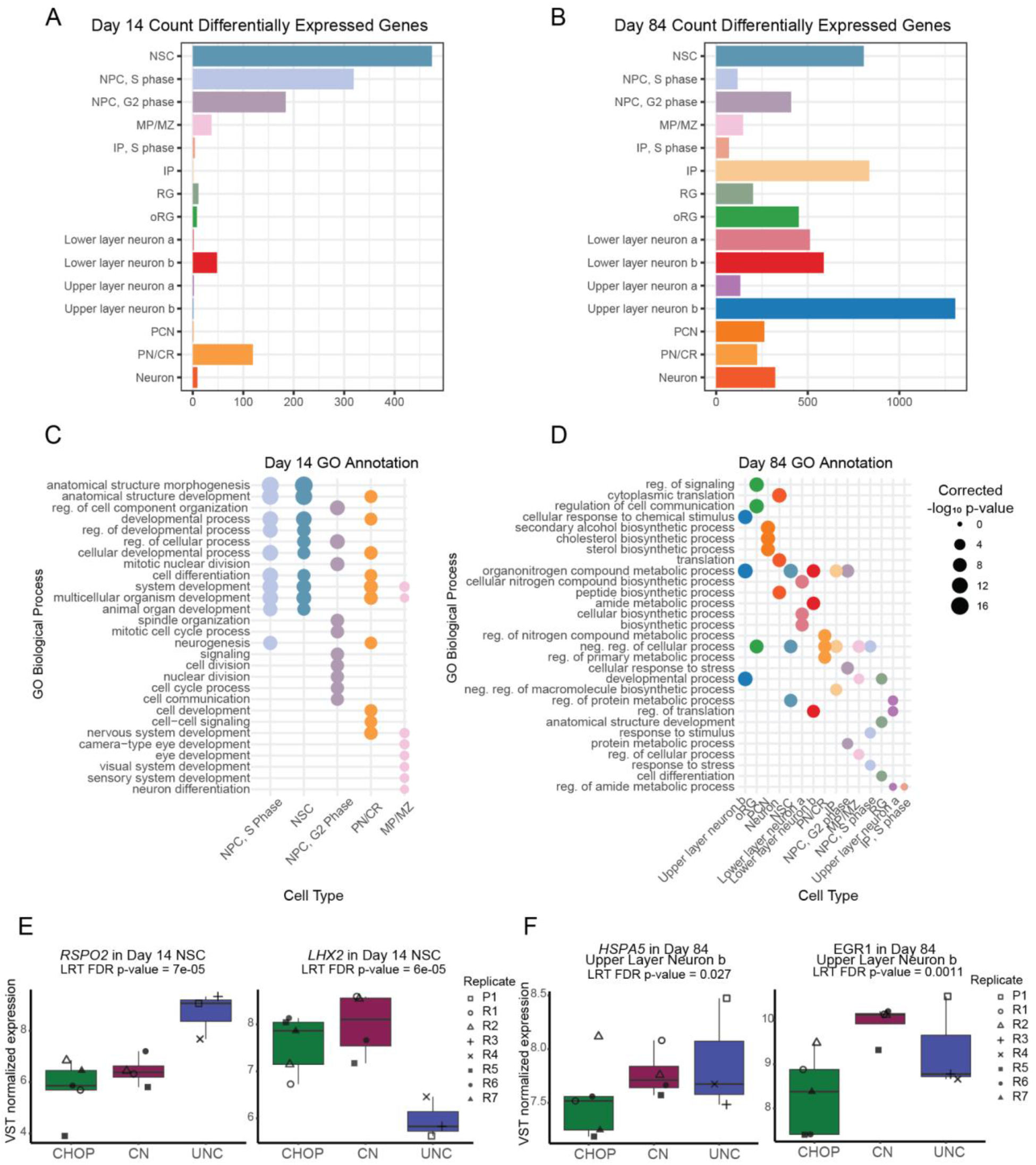
Differential Gene Expression across Sites. Counts of differentially expressed genes in pseudobulked cell types at A) day 14 and B) day 84. Significant gene ontology terms for biological processes associated with differentially expressed genes in cell types at C) day 14 and D) day 84. E) Gene expression differences in *RSPO2* and *LHX2* in day 14 neuroepithelial stem cells (NSC). F) Gene expression differences in *HSPA5* and *EGR1* in day 84 upper layer neuron, group b. Differences in gene expression across-site was determined by likelihood ratio tests (LRT). Abbreviations: regulation (reg), negative (neg).

By day 84, DEGs across sites were present in all cell types and the total number of DEGs increased to 2,188 unique genes across cell types (Figure 3B). DEGs across sites were enriched in metabolic changes and other types of cellular stress, potentially due to differences in necrotic cores and bursting seen across sites (Figure 3D, see also Figure 5H). Examples of DEGs include *HSPA5* and *EGR1*, which had lower expression in CHOP upper layer neurons (Figure 3F). HSPA5 mitigates apoptosis in response to oxygen deprivation in cultured neurons <u>(P. Wang et al. 2015)</u>. *EGR1* upregulation in a perinatal rat model of hypoxia and hypoglycemia resulted in increased proliferation in neural progenitor cells <u>(Alagappan et al. 2013)</u>, suggesting both of these genes promote survival in response to stressors.

Several previous studies have reported cell stress in hCOs with differing interpretations on how impactful the consequences are for modeling cortical development <u>(Bhaduri et al. 2020; Uzquiano et al. 2022; Tanaka et al. 2020)</u>. We additionally used GRUFFI to specifically identify stressed cells independent of cluster annotations based on expression of ER stress and glycolysis associated genes <u>(Vértesy et al. 2022)</u>. Stressed cells were identified only in RG (11.65%) and unspecified neurons (7.06%) and no significant cross-site differences were detected (Supplemental figure 4). This finding is consistent with other groups who have previously observed that RGs and unspecified neurons showed increased ER stress and glycolysis <u>(Uzquiano et al. 2022)</u>, suggesting this kind of cell stress is common across protocols and is not driven by site-specific handling.

Overall, we identified fewer DEGs at day 14, which could be due to greater power to detect gene expression differences at day 84 when more cells were collected for scRNA-seq or an amplification of culture induced technical variation through time. All cell types at day 84 had DEGs, with the majority of significant GO terms implicating cross-site differences in metabolism and cell stress.

### Assaying stem cell state in iPSCs used for hCO differentiation

Rigorous quality controls are recommended for any iPSC differentiation, including validation of undifferentiated cell state by assessing the level of expression of marker genes <u>(Engle, Blaha, and Kleiman 2018; International Society for Stem Cell Research 2023)</u>. To assess stem cell state, gene and protein expression were measured using qPCR and flow cytometry in the same iPSC cultures that went into downstream differentiations (Figure 4). Across sites all iPSC cultures were in an undifferentiated state marked by expression of *OCT4*, *NANOG*, and SSEA3/4. CHOP had higher expression than either other site for *NANOG*, but this did not survive correction for multiple comparisons. Though UNC showed increased variability in SSEA3/SSEA4 populations, leading to nominally significant cross-site differences, all sites had greater than 85% expression of this marker.

**Figure 4.**
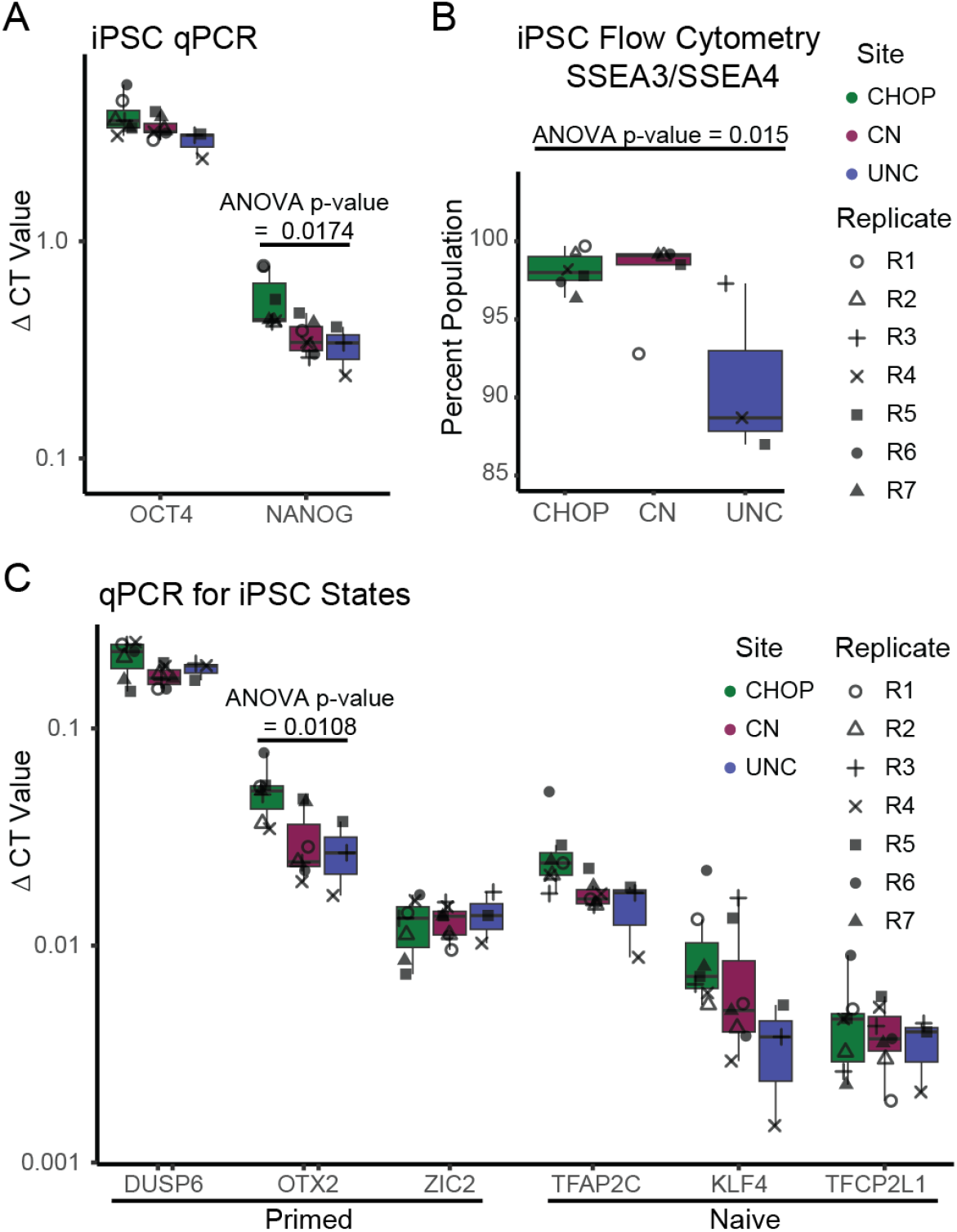
Evaluation of Marker Expression at the Pluripotent Stage and Day 35 across Sites. A) qPCR from iPSCs directly before organoid differentiation for pluripotency. B) Flow cytometry for the pluripotency marker SSEA3/SSEA4. C) qPCR for markers of primed and naive cell state. p-values from one-way ANOVA across site differences are shown when nominally significant. No cross-site differences were detected when correcting for multiple comparisons.

**Figure 5.**
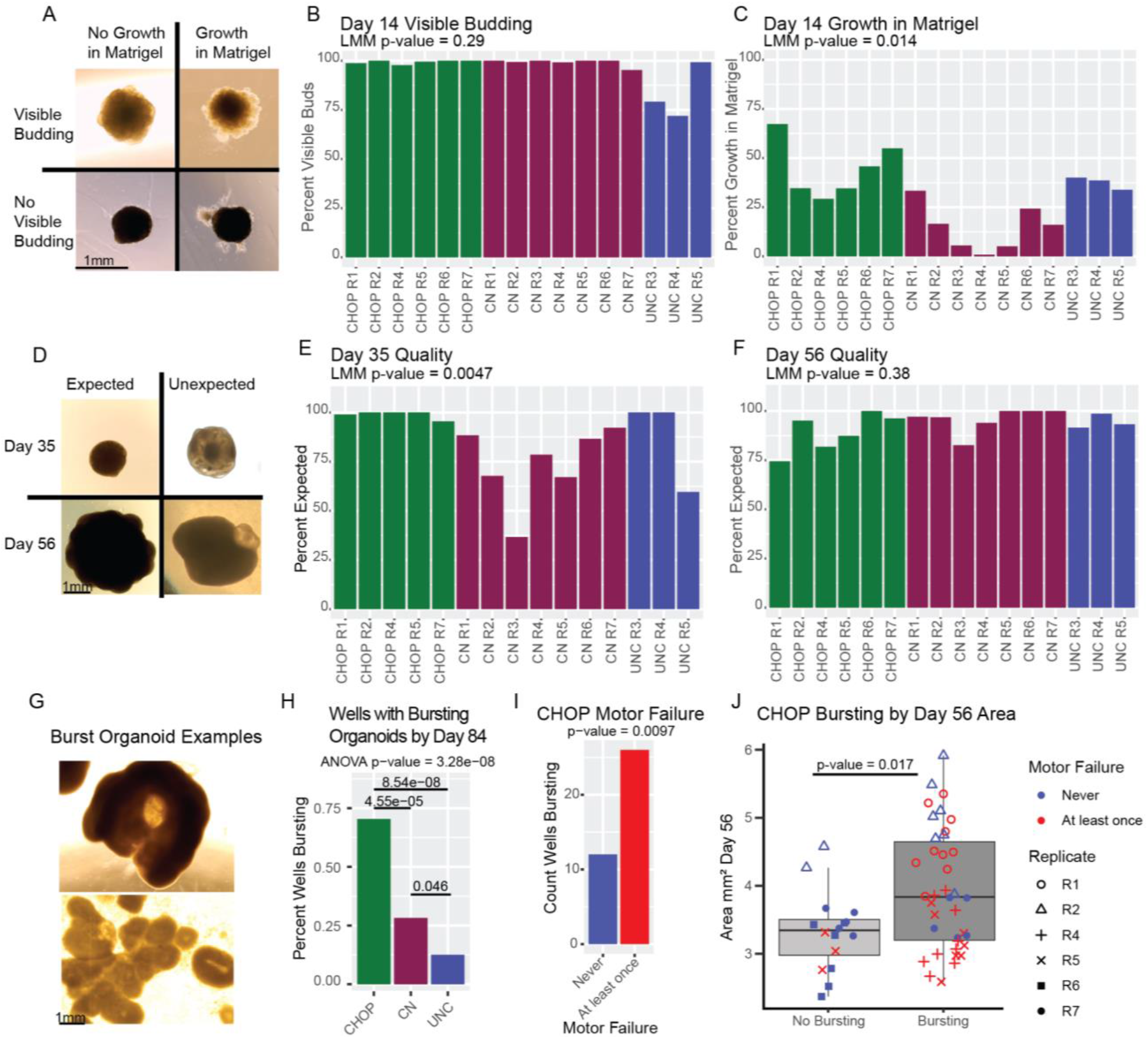
Qualitative Assessments of Organoid Differentiations. A) Representative images of ranking embedded organoids at D14. B) Percent organoids with visible budding at day 14. C) Percent organoids with growth into the matrigel at day 14. D) Representative images of ranking organoids in the bioreactor at day 35 and day 56. E) Percent organoids with expected morphology at day 35. F) Percent organoids with expected morphology at day 56. Significance of cross site differences was evaluated with an ANOVA implemented in a linear mixed effect logistic regression model controlling for the non-independence of multiple organoids within the same differentiation batch using a random effect, and controlling for the ranker who evaluated the images with a fixed effect. G) Example images or burst organoids from CHOP at day 84. H) Percent of wells with burst organoids across sites. I) Count of wells with burst organoids by motor failure at CHOP. J) Organoid day 56 cross-sectional area and motor failure in CHOP burst organoids. Significance of motor failure and cross-sectional day 56 organoid area was evaluated in a linear mixed effect logistic regression model controlling for the other factor.

We also evaluated markers of naive or primed cell pluripotent stem cells <u>(Messmer et al. 2019; Collier et al. 2017; Takahashi, Kobayashi, and Hiratani 2018)</u>. The iPSCs exist along a continuum of transcriptomic states that resemble naive or primed stem cells <u>(Messmer et al. 2019)</u>. Naive stem cells, which resemble the pre-implantation embryo, are hypothesized to be more labile and capable of differentiating to any cell type, whereas primed stem cells, which resemble the post-implantation embryo, are thought to already be directed toward a differentiation fate <u>(Weinberger et al. 2016)</u>. We did not observe cross-site differences in naive or primed markers, other than a nominal difference in *OTX2* (ANOVA adjusted p-value = 0.069). Overall, we detected limited cross-site differences in stem cell gene expression by flow cytometry or qPCR.

### Qualitative ranking of organoids varies by site

hCOs were pre-selected for scRNA-seq based on explicit qualitative criteria (Supplemental Figure 2), so are not representative of all hCOs within the differentiation replicate. Qualitative assessment of organoid morphology using phase contrast microscopy is common in organoid protocols as a way of quickly and cheaply determining the success of organoid differentiation <u>(Qian et al. 2018; Lancaster et al. 2017; Paşca et al. 2015)</u>. Neuroepithelial buds are a model of the developing cortical wall and are a marker of successful telencephalic differentiation (Figure 5A). Growth into the matrigel can indicate off-target differentiation or neurite outgrowth <u>(Lancaster et al. 2013)</u>. To determine the reproducibility of visually assessed organoid quality, we imaged 80 to 100 hCOs per differentiation replicate at day 14 and visually classified each image (Figure 5B-C). We did not observe significant differences across sites for visible budding. However, we did observe cross-site differences for growth into matrigel (p-value=0.014).

At later differentiation time points when hCOs are in the bioreactor, we expect smooth edges with visible budding, but some organoids also have unexpected transparent cysts or thinner strings of cells around the organoid <u>(Qian et al. 2018)</u>. We visually classified 60 - 75 hCOs per differentiation replicate with expected and unexpected morphology at day 35 and 56 (Figure 5D-F). Cross-site differences in organoid morphology were detected at day 35 (p-value = 0.0047). However, by day 56, significant differences were no longer detected. The majority of hCOs with unexpected morphology at day 56 had regions visually similar to choroid plexus <u>(Pellegrini et al. 2020)</u> which could be due to increased medialization caused by WNT stimulation in the protocol. Qualitative assessment using phase contrast microscopy may be helpful for monitoring the differentiation and selecting hCOs for other assays, but hCO morphology across all organoids within a differentiation replicate is inconsistent across sites.

Unexpectedly, after day 56 some hCOs burst and expelled their necrotic cores, leading to dead cells in the well, flattened sheets of neuroepithelial buds, and hollowed cup-like hCOs that can continue to grow (Figure 5G). CHOP had more burst hCOs than the other sites (Figure 5H). Cross-sectional area at day 56 was associated with an increased incidence of bursting when statistically controlling for motor failure (where the motor slowed and stopped spinning) and vice versa, at CHOP (Figure 5I-J). Bursting first occurred within a few days of motor failure. We speculate that CHOP’s hCOs had an increased necrotic core size earlier in the differentiation, due to less diffusion of media through the larger hCOs. Loss of agitation from the motor failure could also exacerbate impaired media diffusion and increase necrotic core size.

### Neuroepithelial bud structure in hCOs

We evaluated intact hCO structure using iDISCO+ tissue clearing <u>(Renier et al. 2016)</u>, immunolabeling, and fluorescent microscopy. We attempted imaging 3 hCOs per differentiation replicate per time point. Neuroepithelial bud lumen size has been used to measure neuroepithelial bud expansion, which is influenced by progenitor packing and proliferation rate <u>(Benito-Kwiecinski et al. 2021)</u>. All sites generated hCOs with neuroepithelial buds, labeled with NCAD+ lumens and PAX6+ ventricular-zone-like (VZ-like) regions at day 14, modeling the developing cortical wall (Figure 6A, Supplemental video 1). Quantification of immunolabeled voxels showed cross-site differences in organoid volume (FDR corrected p-value = 0.0032, Figure 6b), but when normalizing measures of structure for overall organoid volume we did not detect cross-site differences in measures of neuroepithelial bud size (Figure 6C,D). Lumen expansion did not significantly differ across sites when correcting for multiple comparisons, either when NCAD+ area was normalized relative to PAX6+ volumes, representing cortical VZ-like regions only, or when NCAD+ area was normalized to full hCO volumes, representing buds with either PAX6+ RG or PAX6-regions that are likely NSCs.

**Figure 6.**
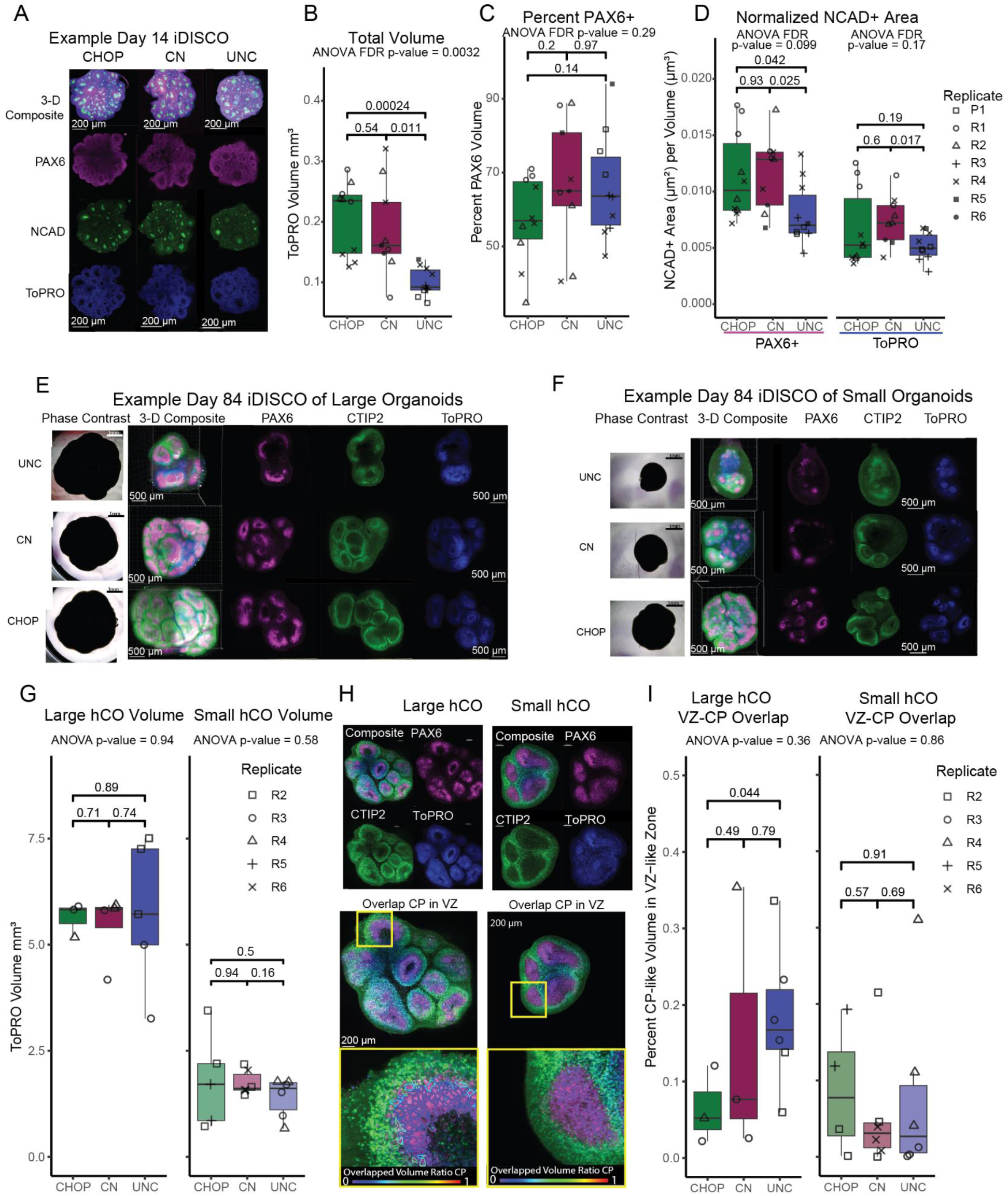
3D Structures in Organoids at Day 14 and Day 84. A) Example image of day 14 organoids from each site. All scale bars are 200 μm except for UNC. B) Total volume of day 14 organoids across sites. Quantification of neuroepithelial bud structure at day 14 as C) percent PAX6+ volume per organoid and D) total NCAD+ area normalized to PAX6+ volume and TOPRO+ volume. Examples images of E) largest organoids and F) smallest organoids from each site at day 84. All scale bars are 500 μm. G) Total volume of largest and smallest organoids across sites. H) Representative 3-D image of PAX6 and CTIP2 with annotated ventricular-zone-like-volumes (VZ-like) and annotated cortical-plate-like-volumes (CP). I) Quantification of volume overlapped between VZ-like and CP-like annotations. All scale bars are 200 μm.

By day 84 neuroepithelial buds had developed into cortical wall-like structures with dense PAX6+ VZ-like regions and dense CTIP2+ cortical plate-like (CP-like) regions (Figure 6E,F, Supplemental video 2). Additionally, all sites had a range of hCO volumes. We attempted imaging the largest and smallest hCO for each differentiation replicate to determine if organoid architecture varied by hCO size. There were significant differences in size between the large and small hCOs for every site, as expected (Student’s t-test, p-value < 0.004 for each site). No cross site differences in total volumes were detected in either large or small hCOs, potentially because the largest hCOs from other sites had burst prior to collection (Figure 6G, Figure 5G-J). Dense VZ-like regions and CP-like regions were semi-automatically annotated to measure their overlapping volumes, where we expect VZ-like regions should be well separated from CP-like regions in high quality differentiations (Figure 6H). All hCOs had less than 40% overlapping volumes, and no significant cross-site differences were detected in small and large hCOs (Figure 6I). Our data support separated cortical wall-like organization in hCOs regardless of size, suggesting internal organization is independent of size. In total, our data suggests that organoid volume was not reproducible across site, at least at day 14, but reproducible measures of organoid structure could be obtained after controlling for overall organoid size.

### Primed markers prior to differentiation predict cell type proportions

In order to find factors that led to cross-site differences or that are predictive of future hCO quality, we tested whether technical variables or assays at earlier time points correlated to cell type proportions measured in sc-RNAseq at later time points. In day 84 organoids, passage at seed (r = 0.82, adjusted p-value = 0.021) and the percent of hCOs with visible buds at day 14 (r = −0.95, adjusted p-value = 0.00012) both correlated to pan cortical neuron proportions (Figure 7A). The effect was driven by two outlier UNC differentiation replicates (Supplemental Figure 6A-D), so it is not possible to disambiguate if handling differences across site or these specific factors resulted in the differences in cell-type proportions.

**Figure 7.**
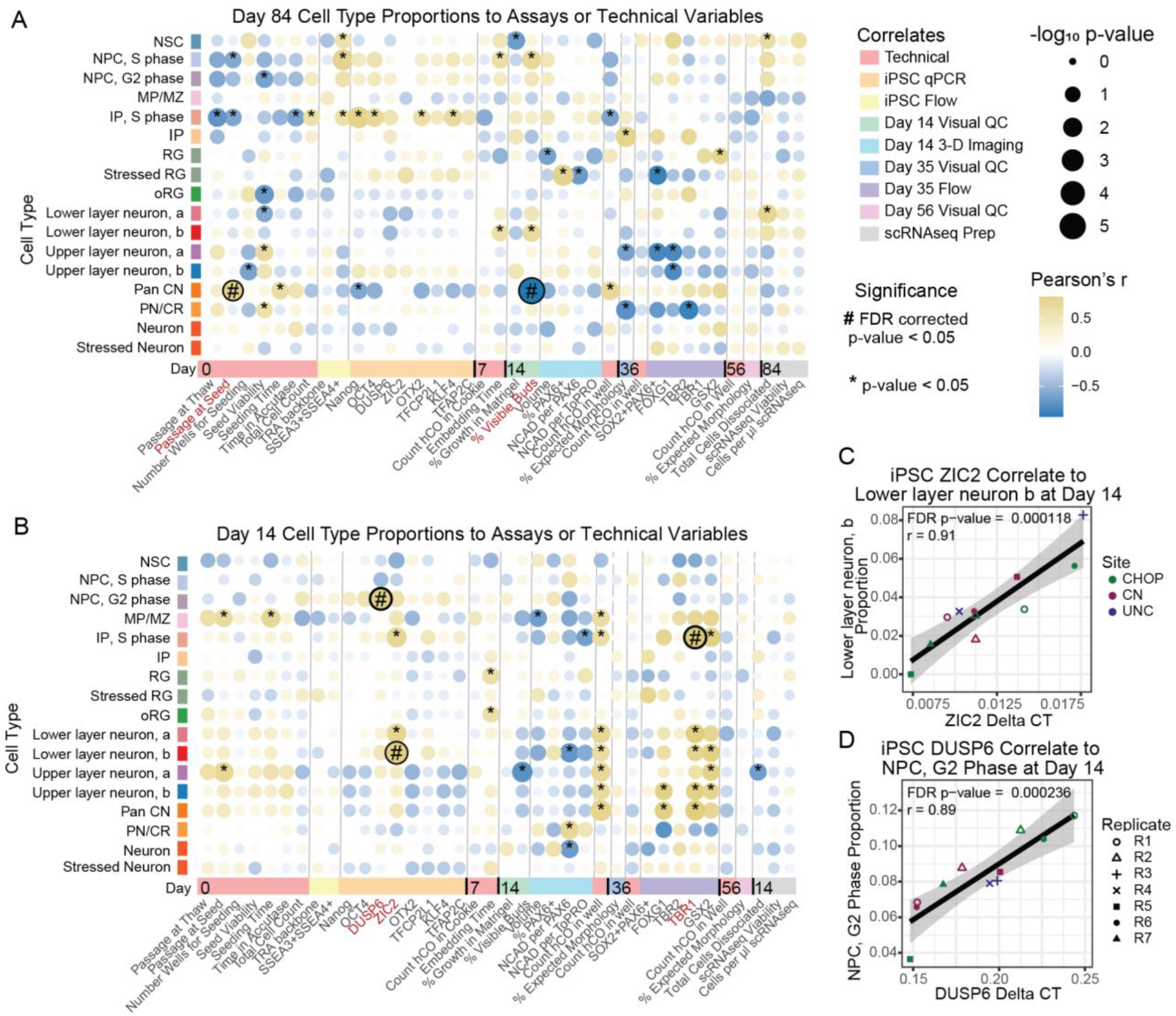
Correlations to Cell Type Proportions. Pearson’s correlations of cell type proportions in scRNAseq data to assay results or technical variables at A) day 84 or B) day 14. C) Correlation between primed marker ZIC2 and lower layer neuron, b proportions at day 14. D) Correlation between primed marker DUSP6 and NPC, G2 phase in day 14 organoids.

Two primed stem cell state markers, which are highly correlated to each other (r = 0.75), both predicted cell type proportions at day 14. *ZIC2* expression is positively correlated with lower-layer neuron group b (r = 0.91, adjusted p-value = 0.000118). *DUSP6* expression correlated to increased proportions of NPCs in G2-phase (r= 0.89, adjusted p-value = 0.00024; Figure 7b). Importantly, no individual site is an outlier driving these results (Figure 7C-D). This suggests that the pluripotency state can influence downstream differentiation potential, at least at day 14. The proportion of IPs in S-phase was also correlated with the TBR1+ population measured by flow cytometry at day 35. Though the proportion of both cell populations is small (1.57% IPs in S-phase, 4.0% TBR1+ populations), it is expected that proliferating IPs will lead to an increase in the total number of neurons <u>(Rakic 2009)</u>. Identification of an expected relationship between progenitor and neuronal cell types across assays increases the confidence in our analysis. Our results suggest iPSC transcriptional state influences downstream differentiation outcomes.

## Discussion

Our study aimed to identify hCO phenotypes that are reproducible across sites using harmonized differentiation protocols. We adapted the original protocol to feeder-free iPSC culturing conditions and generated embryoid bodies from single cell suspensions, which necessitated increased concentration of dual SMAD-inhibitors to yield similar cell type production later in the differentiation <u>(Qian et al. 2016, 2018)</u>. Like the original protocol, hCOs generated across three sites from one iPSC line produced similar proportions of mostly cortical cell types (Figures 1 and 2). All sites generated hCOs with similar neuroepithelial bud architecture mimicking the cortical wall (Figure 6). We detected cell-type specific differentially expressed genes across sites as early as day 14 of the differentiation (Figure 3). Interestingly, genes involved in cellular stress and metabolism were differentially expressed later as organoids grew in size (Figures 3 and Figure 6). Non-reproducible measures across sites included organoid size, hCO morphology, and bursting (Figure 5). We also found that variations in primed stem cell markers prior to differentiation correlated with proportions of cortical cell types in day 14 hCOs, suggesting that small changes in stem cell state can influence cell type proportions in differentiated hCOs (Figure 7). Overall, our data produced promising results, supporting reproducibility of cell type proportions and structure between the sites and across replicates, suggesting these measures may be useful for prospective cross-site meta-analysis future studies.

### Reproducible cell type proportions and different transcriptomes

Irreproducibility of cell type composition was cited as a major barrier to using hCOs in high-throughput drug screening and modeling neurodevelopmental disorders <u>(Huch et al. 2017)</u>. Unguided organoid protocols have produced an inconsistent variety of non-cortical cell types and cell type composition across organoids <u>(Lancaster et al. 2013; Quadrato et al. 2017)</u>. In contrast, guided protocols show more consistent cell-type composition within a given lab and across multiple iPSC lines with minimal non-cortical cells <u>(Velasco et al. 2019; Yoon et al. 2019)</u>. Here, we extend these findings to show that despite cross-site differences in handling, cell-type composition in a guided differentiation protocol is consistent and mainly cortical. Though we showed reproducibility in cell-type proportions, we also recapitulated known limitations in the fidelity of hCO models. All sites had reproducible non-cortical differentiation, had glycolytic and ER stressed cells, and broad neuronal cell types that do not closely resemble a particular cortical layer or region <u>(Tanaka et al. 2020; Bhaduri et al. 2020; Kelley and Pașca 2022)</u>. These known limitations could be accounted for in future experimental designs by bioinformatically filtering out stressed cells or cells that do not model the brain region of interest <u>(Vértesy et al. 2022)</u>. Discrepancies were noted between transcripts detected in scRNAseq and protein levels detected by immunolabeling in flow cytometry and 3D imaging, which may be due to a number of factors including insufficient read depth <u>(Pokhilko et al. 2021)</u>, organoid selection, and/or differences between the half-life of transcripts and corresponding proteins <u>(Edfors et al. 2016)</u>.

Differentially expressed genes implicated metabolic and cellular stress differences across sites in several cell types. Given that we observed volume differences in hCOs across sites, and increased cross-sectional area was associated with bursting organoids at one site, differential expression of stress and metabolism genes could be due to lack of media diffusion through the organoid and large interior necrotic cores. Using a guided hCO protocol that minimizes necrotic cores <u>(Watanabe et al. 2017; Qian et al. 2020)</u> or protocols that incorporate vascularization into hCO generation <u>(Cakir et al. 2019)</u> could reduce the possibility of these metabolic-related cross-site differences.

### Cortical wall-like organization across time and size

We observed consistent neuroepithelial budding at day 14 leading to cortical wall-like organization at day 84 hCOs, regardless of size. This result demonstrates that a key feature of hCO culture is reproducible across sites. Cortical wall-like organization increases cell-to-cell contact in hCOs, promoting oRG and IP production <u>(Scuderi et al. 2021)</u>, and enables the study of defects in neuronal migration <u>(Blair, Hockemeyer, and Bateup 2018)</u>. Though not significantly different across sites, some organoid structural phenotypes were still variable within sites which could be due to technical factors like variability in clearing, antibody penetration, or agarose embedding distorting the light path in addition to biological influences. Tissue clearing and analysis are still developing and are not yet standardized in hCO studies, though progress is being made in developing analysis tools for organoid structures in 3D that could increase reproducibility in the future <u>(Benito-Kwiecinski et al. 2021; Albanese et al. 2020; Y. Li et al. 2017)</u> hCO size varies across site and within each differentiation. Size was not previously addressed in single site studies of hCO reproducibility <u>(Velasco et al. 2019; Yoon et al. 2019)</u>. However, differences in hCO size in case-control comparisons are consistently documented when large effects are expected, such as when studying Zika virus <u>(Watanabe et al. 2017; Qian et al. 2016)</u>, monogenic causes of macrocephaly <u>(Y. Li et al. 2017; Zhang et al. 2020; Urresti et al. 2021)</u> and microcephaly <u>(R. Li et al. 2017; Lancaster et al. 2013)</u>. Though large effect genetic variants or viral infection may overcome the noise in hCO size measurements at a single site, our study suggests that hCO size is not suitable for cross-site meta-analysis using the current differentiation protocol. Instead hCO size and morphology are important technical factors to report and statistically control.

### iPSC cell state influences differentiation

A previous study using different iPSC culturing conditions to modulate stem cell state found that an intermediate pluripotency state between naive and primed led to higher quality hCOs with fewer non-cortical cell types and more robust cortical wall-like organization <u>(Watanabe et al. 2022)</u>. We found that even with culturing the same iPSC line under the same conditions, increased expression of primed marker *ZIC2* prior to differentiation led to a greater proportion of lower layer cortical neuron production, and increased *DUSP6* expression led to increased proliferating progenitors early in the differentiation. ZIC2 is known to regulate gastrulation and dorsal forebrain specification <u>(Houtmeyers et al. 2013)</u>. DUSP6 has been implicated in cancer and regulates the MAPK pathway <u>(Ahmad et al. 2018)</u>. We did not observe a significant association with naive or primed gene markers and non-cortical cell types or unspecified neurons. While not directly evaluating the same phenotypes <u>(Watanabe et al. 2022)</u>, these results suggest that stem cell state during pluripotency may impact hCO phenotypes after differentiation. We recommend that future studies monitor variation in iPSC cell state before hCO differentiations <u>(International Society for Stem Cell Research 2023)</u>. Though primed and naive markers are currently not standard for quantitative iPSC quality controls, like the qPCR score card <u>(Fergus, Quintanilla, and Lakshmipathy 2016)</u>, gaining an understanding of how stem cell state influences multiple hCO protocols could lead to better standardization of iPSC quality and more reliable hCO phenotypes.

### Limitations

Our study evaluated cross-site hCO reproducibility across many phenotypes using one cell-line (PGP1) and one differentiation protocol. As such, we are not able to assess variability driven by differences in cell lines or protocol. Line-to-line variability may also influence differentiation outcomes, caused by either genetic differences or technical variation in iPSC generation. Recent work has shown that variability in iPSCs is driven by both genetic and technical variation, though this has not been evaluated in hCOs derived from many donor iPSCs <u>(Kilpinen et al. 2017; Burrows et al. 2016)</u>. Secondly, differences in cell stress related genes may not apply to protocols without necrotic cores, and differences in WNT related genes may not be relevant to protocols not activating WNT during the differentiation. To demonstrate hCO reproducibility in future studies, we recommend using the same iPSC line cultured across sites with the same protocol as a “rosetta” line <u>(Volpato and Webber 2020)</u>. Finally, though we measured multiple phenotypes from multiple differentiation replicates, cross-site differences may exist that we were underpowered to detect for all measurements used here.

## Conclusions

Several fields have undergone a reproducibility crisis <u>(Marek et al. 2022; Border et al. 2019; Baker 2016; Open Science Collaboration 2015)</u>. Fields that have overcome this reproducibility crisis were able to do so by: 1) demonstrating reproducibility of measurements, 2) standardizing data acquisition and sharing and 3) combining data from multiple sites to increase statistical power <u>(Medland et al. 2022; Sullivan et al. 2018; Moshontz et al. 2018)</u>. hCO studies of neuropsychiatric disorders are in their infancy and may expect similar reproducibility problems unless proactively addressed. This study demonstrates that hCOs can generate reproducible cell types and organization when produced across multiple sites, suggesting that comparisons of changes in cell type proportions or organoid cortical wall like organizations can be properly controlled, enabling future prospective meta-analytic iPSC-derived hCO studies in neuropsychiatric disorders.

## Methods

### iPSC Culture and Maintenance

PGP1 vials of the same passage were shipped to each site from Children’s National. PGP1 was thawed at each site and maintained on hESC-qualified matrigel (Corning 354277) coated plates in mTeSR+ (Stem Cell Technologies 100-0276) and fed every day except for UNC pilot one and the first week of UNC replicate 3, which was fed every other day, as described in the mTeSR+ user’s manual. Cells were passaged when 80% confluent using Versene (Thermo Fisher 15040066) or 0.5mM EDTA in DPBS. Mycoplasma testing using PCR detection was performed before each experiment and no mycoplasma was detected at any site. PGP1 was karyotyped at CN before the cell line was distributed to all the sites and no chromosomal abnormalities were detected.

### Organoid Protocol

The protocols for organoid differentiation and collection of organoids for each assay were harmonized across sites. This included practicing setting up the bioreactor, the first month of the differentiation, and the single cell dissociations for flow cytometry, and the visual quality of organoids going into the scRNA-seq experiment. Additionally, a minimum number of organoids at each step of the differentiation was agreed upon, for example at least 92 organoids were needed at day 14 to proceed with a differentiation replicate. Due to concerns about lot-to-lot variability in matrigel, the same lot of matrigel was used across three sites, except for UNC pilot 1. All other culturing reagent lots were not standardized. Culture plates and bioreactors were the same across the sites. All sites performed differentiation replicates in parallel and started with PGP1 between passages 38-42. Each site attempted the differentiation at least once prior to including any data in this study in order to harmonize the protocol, standardize and practice harvesting procedures for downstream assays, and reach agreement on the technical variables to record.

The organoid differentiation protocol was adapted from (Qian et al. 2018) to feeder-free conditions. Briefly, the major changes included culturing the iPSCs on hESC qualified matrigel in mTeSR+, generating embryoid bodies from a single cell suspension (Eiraku et al. 2008), and doubling the concentrations of Forebrain First media small molecules (described below). In more detail, after at least 2 passages of the iPSC line, cultures were harvested with Accutase (Thermo Fisher A1110501), centrifuged and resuspended in recovery media (6 μM Y27632 (Fisher Scientific ACS-3030) in mTeSR+), and seeded at nine thousand cells per well into a 96 well v-bottomed ultra-low attachment plate (S-Bio MS-9096VZ) in recovery media (6 μM Y27632 (Fisher Scientific ACS-3030) in mTeSR+). The following day, half of the media was replaced with 2X Forebrain First Media (DMEM/F12 Thermo Fisher 11320033, 20% Knock-Out Replacement Serum Life Technologies 17502048, Glutamax Life Technologies 35050-061, MEM-Non Essential Amino Acids Gibco 11140050, Pen-Strep Gibco 15140122, Beta-Mercaptoethanol Thermo Fisher Scientific 21985023) with double the concentration of dual-SMAD inhibitors (8uM Dorsomorphin Sigma-Aldrich P5499, 8uM of A-83 STEMCELL Technologies 72022), initiating neural induction and termed Day 0. From Day 1 to Day 4, three fourths of the media was replaced with Forebrain First Media with dual SMAD inhibitors (4uM Dorsomorphin, 4uM A-83). Day 6 and Day 7, half of the media was replaced with Forebrain Second Media (DMEM/F12 Thermo Fisher 11320033, N2 Life Technologies 17502048, Glutamax Life Technologies 35050-061, MEM-Non Essential Amino Acids Gibco 11140050, Pen-Strep Gibco 15140122, Beta-Mercaptoethanol Thermo Fisher Scientific 21985023) with a SMAD inhibitor (SB-431542 Fisher Scientific 16-141) and WNT activator (CHIR-99021 Fisher Scientific S12632). At Day 7, organoids were removed from the 96 well plate and embedded in a 60% Growth Factor Reduced Matrigel (Corning 356230) and Forebrain Second mixture. Briefly, the growth factor reduced matrigel was diluted to 60% in Forebrain Second mixture and mixed with organoids, then plated on to ultra-low attachment six well plates (Sigma-Aldrich CLS3471) and allowed to solidify at 37° C for 30 minutes, followed by the addition of 3 mL of Forebrain Second media per well. From days 8 to 13 all media was replaced with fresh Forebrain Second media every other day.

At Day 14 organoids were extracted from the matrigel with gentle pipetting and ∼10 organoids were placed into each well of non-tissue culture treated 12 well plates with individual wells from the matrigel step kept separate from other wells. Each well was fed with 4 mL of Forebrain Third Media (DMEM/F12 Thermo Fisher 11320033, N2 Life Technologies 17502048, Glutamax Life Technologies 35050-061, MEM-Non Essential Amino Acids Gibco 11140050, Pen-Strep Gibco 15140122, Beta-Mercaptoethanol Thermo Fisher Scientific 21985023, B-27 Life Technologies 17504044, 2.5ug/mL Insulin Sigma-Aldrich I9278-5ML). Lids of the 12 well plates were replaced with miniaturized spinning bioreactor (SpinΩ bioreactor). The DC motor was set to 7.5 V or ∼100 rotations per minute of the internal paddles. Organoids were fed every 2 to 3 days by entirely replacing the media with fresh Forebrain Third Media. Starting on Day 35 growth factor reduced matrigel (1:100) was supplemented into cold F3 media and warmed to room temperature before feeding. If the media was turning yellow (evidence of increased acidity) or the organoids were developing cysts, the organoids were separated out into new wells. From Day 70 to Day 84, organoids were fed with Forebrain Forth Media (Neurobasal Life Technologies 21103-049, Ascorbic Acid Sigma-Aldrich A4544, Glutamax Life Technologies 35050-061, MEM-Non Essential Amino Acids Gibco 11140050, Pen-Strep Gibco 15140122, B-27 Life Technologies 17504044, cyclic AMP 0.5 mM Sigma-Aldrich A6885, 20 ng/mL BDNF Life Technologies PHC7074CS1, 20 ng/mL GDNF R&D Systems 212-GD-010). Debris from burst organoids were removed from wells, when possible, with attempts made to retain remaining viable organoids.

### qPCR for iPSC cell state

300,000 iPSCs were harvested on Day −1 from extra iPSCs not used for seeding the organoids. Total RNA was extracted from cells using the Trizol Plus RNA Purification Kit (Thermo Fisher Scientific,12183555) according to the manufacturer’s instructions. The Prime Script RT Master Mix (Takara Bio, RR036A) was used to synthesize cDNA according to the manufacturer’s instructions, and qRT-PCR was performed on the QuantStudio 7 Flex Real-time PCR System (Thermo Fisher Scientific) with TB Green Premix Ex Taq Ⅱ (Takara Bio, RR820B). Primers are in the supplemental materials. Delta CT values relative to the housekeeping gene EIF4A2 are reported for each gene because there is no control group. One-way ANOVAs were used to test for cross-site differences with BH-FDR. Post-hoc testing with student-t tests were used to test for which site was different from the others.

### Phase Contrast Imaging & Analysis

At days 0, 7, 14, 35, 56, 70 and 84, phase contrast images were acquired of the organoids on an EVOS XL Core microscope using the 4x objective. CN acquired phase contrast images with Olympus CKX31 through a LabCam lens. Due to non-rigid image distortions from differences in image acquisition, phase contrast images were used for qualitative assessment only and could not effectively be evaluated for cross-site size comparisons. A custom ranking scale for organoids was designed for Day 14, Day 36 and Day 56 based on visual morphology of the organoids. Rank scales used for training are in supplemental materials. Three independent rankers were trained on the rank scale until 85% inter-rater agreement was reached on a test set. Cross site differences were evaluated with an ANOVA implemented in a linear mixed effect logistic regression model using a random effect to control for the non-independence of multiple organoids within the same differentiation batch, and controlling for the ranker who evaluated the images with a fixed effect.

### Tissue Clearing and Imaging

Organoids for imaging were fixed in 4% paraformaldehyde (PFA) for 30 minutes at CN and UNC, and for 1 hour at CHOP. Day 14 and Day 84 organoids were tissue cleared using an adapted iDISCO+ protocol (Renier et al. 2014). In microcentrifuge tubes, hCOs were washed in phosphate-buffered-saline (PBS), permeabilized for at least 15 hours (20% dimethyl-sulfoxide, Fisher, BP2311; 1.6% Triton X-100, Sigma-Aldrich, T8787; 23 mg/mL Glycine, Sigma-Aldrich G7126), and incubated in blocking buffer 6% goat serum (Abcam, ab7481) for at least 2 hours. hCOs were then transferred to individual wells of a 96 well plate and incubated with antibodies for 1 day for day 14 hCOs or 3 days for day 84 hCOs at 37° C in PTwH buffer (PBS; 0.5% Tween-20, Fisher, BP337; 10 mg/L Heparin, Sigma-Aldrich, H3393) while on an orbital shaker at 90 r.p.m. After 3-5 washed with PTwH, hCOs were then incubated with TO-PRO-3 and secondary antibodies for 1-3 days at 37° C with agitation (Supplemental resource table 3).

Samples were then washed 3-5 with PTwH and then embedded in 1% agarose blocks between 2-5mm thick. Agarose blocks of organoids were dehydrated in 5mL tubes using a graded methanol series of 30 minute incubations (Fisher, A412SK), incubated in 66% dichloromethane/methanol (Sigma-Aldrich, 270997) for 1.5 hours and washed in 100% dichloromethane twice before being stored in a dibenzyl ether solution (RI = 1.56, Sigma-Aldrich, 108014). Samples were imaged within 3 days of completing the tissue clearing.

Day 14 organoids (<1mm in diameter) were imaged on an Olympus FV3000RS, using an Olympus LUCPlanFL-N 20x long working distance objective and Galvano scanning. Resolution was 0.621 x 0.621 x 3.5 μm/voxel. 3-5 organoids were imaged per replicate, and each agarose block contained one replicate from each site to reduce batch effects.

Day 84 organoids (3-5 mm in diameter) were imaged using Imspector software on a LaVision Ultramicroscope II light sheet microscope with a 2X/0.5NA MVPLAPO objective and CMOS camera (Andor). Resolution was 0.95 x 0.95 x 4.5 μm/voxel. Each experiment was in a unique agarose block and each organoid was imaged in 1 to 4 tiles. Images were taken with either a single or multiple light sheets and with or without dynamic horizontal focusing, depending upon image quality.

Raw files were converted into IMARIS files using ImarisFileConverter (v9.9.0 or 9.7.0). For multi-tile acquisitions, IMARISStitcher (9.10.0 or 9.7.0) where images were placed on a grid, aligned and stitched. IMARIS 10.0 was used to create surface annotations for day 14 and day 84 images. All images were first batch processed following auto adjustment of intensity across all channels. Batch processing used automatic intensity thresholds and intensity based splitting with 20 micron seed points to create surfaces. An independent experimenter reviewed these batch processed images. Adjustments to the surfaces were made by changing the intensity threshold, the intensity base splitting, manually removing actions of the surfaces, filtering by the split surfaces by total volume or mean-intensity of other channels in the surfaces to account for potential autofluorescence and intensity differences across tiles. For day 14 hCOs, surfaces were created for ToPRO, NCAD, and PAX6. For day 84 hCOs, surfaces were created for dense PAX6+ ventricular-zone-like volumes, dense CTIP2+ cortical wall-like surfaces, and total organoid volume. “Surface to surface statistics”, which includes calculating total overlapping volumes between CTIP2+ surfaces and PAX6+ surfaces, were calculated in IMARIS. One-way ANOVAs were used to test for cross-site differences with BH-FDR used for multiple comparisons. Post-hoc testing with student-t tests were used to test pairwise differences between sites.

### Flow Cytometry

Flow cytometry samples were harvested by pre-treatment in collagenase (1mg/ml) + DNase (0.01mg/ml) at 37°C (Days 14 and 35 only), followed by trypsin for 5 minutes, then fixation with 1.6% PFA for 30 minutes (days 7, 14 and 35). Immunolabeling was performed by incubation in primary antibody, followed by fluorescent-conjugated secondary antibody (when primary antibody is not directly conjugated), each for 1 hour at room temperature. FACS buffer (Dulbecco’s Phosphate Buffered Saline + 0.5% Bovine Serum Albumin + 0.05% sodium azide) was used as the diluent and wash buffer for extracellular antibody epitopes. 1X Intracellular Staining Perm Wash Buffer (diluted in water from Biolegend, cat no. 421002) was used to permeabilize cells and as the diluent and wash buffer for intracellular epitopes. Staining was followed by reconstitution in FACS buffer and flow cytometry readings on a Cytoflex V5-B5-R3 (Beckman Coulter) and data was analyzed with FlowJo (version 10.8.2) (Supplemental resource table 2). Day −1 samples were harvested with initial EB seeding. Compromised samples, due to fixation or storage issues, were identified with a majority trypan blue + stained cells and were omitted from analysis. For the gating, cell populations were gated based on all cell populations and excluding debris and dead/dying cells, as events identified with low FSC-A and/or SSC-A (see Supplemental Figure 4). Multiple cells and clusters were additionally excluded with FSC-W/FSC-A gating. Stem marker (SSEA3+, SSEA4+, and PODXL+) gating was determined based on the no antibody control. PAX6+ cells and FOXG1+ cells gates were determined based on Day −1 cell immunoreactivity, which is negative for both measures. SOX2+ cells were gated from the SOX2+ population identified at Day −1. TBR1+ and GSX2+ cells were determined based off of Day 7 gating, which is negative for both populations. TBR2+ cells were determined in comparison of each cell population measured against immunoreactivity of a Rabbit IgG isotype control (Cell Signaling, #5472). For all measures, except for TBR2, identical gates were applied to each cell population. Cross-site analyses were limited to day −1 and day 35 samples, due to sample loss of days 7 and 14 samples, resulting from site-specific storage issues. One-way ANOVAs were used to test for cross-site differences at day −1 and day 35 with BH-FDR used for multiple comparisons. Post-hoc testing with Student’s t-tests were used to test for pairwise differences between sites.

### scRNA-seq Library Preparation

Organoid papain dissociation was adapted from a previous protocol using papain (Worthington Biochemical Corporation LK003150) (Velasco et al. 2019). At Day 14, organoids were selected for scRNA-seq that had phase bright budding structures and no residual matrigel. At Day 84, a phase contrast image was taken of each organoid harvest for scRNA-seq, which were selected to have smooth edges and no visible cysts. All scRNA-seq samples had at least 70% viability as determined by Trypan blue staining before proceeding. Single cells were permeabilized and fixed using ParseBiosciences Cell Fixation Kits (SB1001). Frozen aliquots of fixed single cells were stored at −80 C and shipped on dry ice overnight to UNC from CHOP and CN. Barcoding and library preparations were completed using 2 Whole Transcriptome Kits (EC-W01030) from Parse Biosciences at UNC.

### Quality control of scRNA-seq data

Raw data was aligned using Salmon (Srivastava et al. 2020) to a reference transcriptome including spliced+intron reads using Gencode (V40) (Frankish et al. 2021) annotation made with make_splici_txome function in the roe package (https://github.com/COMBINE-lab/roe). Gene expression matrices were loaded into R using *loadFry*() of fishpond (v2.0.1) with snRNA mode which includes the spliced, unspliced and ambiguous mRNA count of the gene. Data was processed using the Seurat R package (version v4.0.3) standard pipeline (Hao et al. 2021). Cells were removed if (1) < 1,000 genes per cell were detected, (2) < 1,500 unique mRNAs (UMIs) were detected per cell, or (3) >=10% of total RNA reads were from mitochondria.

Data was split into unique samples defined as a combination of site, differentiation replicate, and organoid differentiation day (14 or 84). Within sample, we estimated doublets using doubletFinder (v2.0.3) (https://github.com/chris-mcginnis-ucsf/DoubletFinder) (McGinnis, Murrow, and Gartner 2019) with an expected doublet rate of 4.25% and pN=0.25. Optimal pK was set to maximum of the bimodality coefficient distribution. After QC, we obtained 133,980 high quality cells in total. Average genes/cell: 3,868 (min: 1,000, max: 22,008, median: 3,369). Average cells/sample: 4,807 (min: 116, max: 23,143, median: 2,886, total samples: 28). Average UMIs/cell: 12,881 (min: 1,500, max: 689,582, median: 7,820).

### Data Normalization and Clustering

To account for batch effects, data was transformed and normalized using *SCTransform*() (v2), separately for each sample and then integrated to one dataset using the top variable 5,000 genes and two references (UNC R3 at day 14 and day 84). *FindNeighbors*() was applied to the integrated dataset to identify and construct shared nearest neighbors. Cells were then partitioned into distinct clusters by adjusting the resolution to 0.6. Clusters were manually annotated by the significantly upregulated genes identified by Seurat *Find.Markers* and expression of known canonical markers. tSNE projection was chosen to visualize cells with gene expression.

### Correlation to Primary Fetal Tissue

A Seurat object containing 5 fetal cortical samples from 6 to 22 gestational weeks was graciously provided by the authors (Bhaduri et al. 2020). Primary tissue data was reprocessed to with the same cell quality as our organoid data set; cells were included if at least 1,000 genes were detected, at least 1,500 unique mRNAs (UMIs) were detected and no more than 10% of total RNA reads were from mitochondria. The intersection of the 5,000 most variable genes for each data set were used to correlate clusters from the current hCO data set and *in vivo* cell types. Gene expression from both data was normalized using SCT Transform. The Benjamini-Hochberg method was used to correct for multiple comparisons.

### Cell Type Proportions

Cell type proportions were calculated for the dataset generated in this manuscript using Propeller (Phipson et al. 2022), which stabilizes variance estimates for proportions of each cell type and performs testing within an empirical Bayes framework. We used a moderated ANOVA test across the 3 sites followed by Benjamini-Hochberg correction for multiple comparisons across cell-types (Supplemental material). No statistical tests were performed for the UNC day 56 hCO data because only one site was collected. Cell type proportions for the (Polioudakis et al. 2019) dataset were reported in the original manuscripts. Cell type proportions for Bhaduri et al., 2020 were calculated based on the authors’ original annotations and plotted using Propeller.

### Differential Gene Expression

Cells were pseudobulked across within assigned cell type and day by summing read counts using Muscat (1.14.0) (Crowell et al. 2020). Summed read counts were calculated for each pseudobulked cluster. Using DESeq2 (1.40.2) (Love, Huber, and Anders 2014), we compared the fit of a model containing dummy variables marking each site to the fit of the model with only an intercept using likelihood ratio testing to determine if any gene showed cross-site differences. We performed this differential expression analysis for any gene with greater than 5 reads across 3 samples for each pseudobulked cell type at both time points (Supplemental DEG). Multiple test correction within each cell type at each time point was performed using the Benjamini-Hochberg procedure (Benjamini and Hochberg 1995).

### Gene Ontology Annotation

Differentially expressed genes were annotated using gost function in gprofiler2 (Kolberg et al. 2020) and only gene ontologies for biological processes are shown (Raudvere et al. 2019). gSCS was used for multiple comparison testing.

### Identification of Stress Cells

Granular functional filtering (GRUFFI) (Vértesy et al. 2022) was used to identify stressed cells (https://github.com/jn-goe/gruffi). Cells were partitioned into granules (equivalent to Res 70 in Seurat) and then assigned scores for GO terms of ER stress, gliogenesis, and glycolytic process. Default thresholds for these GO terms were used to call “stressed” or “unstressed cells”. Individual cells had previously been assigned to a cell type classification and this was used to determine the proportion of stressed cells in each cell type and sample across time.

### Correlations of Technical Variables and Assays to Cell Type Proportions

Assay results and numerical variables recorded throughout the differentiation that were available for all 3 sites were correlated with cell type proportion variability across differentiation replicates. Technical factors recorded and included were: passage at thaw, passage at seeding, number of wells of iPSCs used for seeding, viability of iPSC at seeding, minutes it took to complete seeding, minutes cells were in Accutase, total cell count of iPSCs at seeding, count of hCOs in each matrigel cookie at day 7, minutes it took to complete embedding hCOs in matrigel cookie, average count of hCOs in each well of the bioreactor at day 14, average count of hCOs in each well of the bioreactor at day 35, average count of hCOs in each well of the bioreactor at day 56, total cells dissociated in papain, viability of cells after papain dissociation, concentration of cells for loading the scRNA-seq library preparation. UNC pilot 1 samples did not have most of the technical variables or assay results as the other replicates. The Benjamini-Hochberg method was used to correct for multiple comparisons for day 84 and day 14 separately.

## Supporting information

Supplemental DEG

## Acknowledgements

We thank the Intellectual and Developmental Disabilities Research Center organoid working group for support and insightful discussions. This work was supported by the NIH (3-P50-HD103573-03S1, R01 MH130441, R01MH121433, R01MH12012, R01MH118349), the Foundation of Hope, UNC TraCS (550KR262111), and the UNC School of Medicine to J.L.S, (R01AA025215, R01AA026272) to K.H.T., and F31MH124427-01A1 to M.G. The University of North Carolina High Throughput Sequencing Facility is supported by the University Cancer Research Fund, Comprehensive Cancer Center Core Support grant (P30-CA016086), and UNC Center for Mental Health and Susceptibility grant (P30-ES010126). We thank Pablo Ariel at the Microscopy Service Laboratory and Michelle Itano at the Neuroscience Microscopy Core for assistance with imaging. The Microscopy Services Laboratory, is supported in part by P30 CA016086 Cancer Center Core Support Grant to the UNC Lineberger Comprehensive Cancer Center and the North Carolina Biotech Center Institutional Support Grant 2016-IDG-1016. The University of North Carolina’s Neuroscience Microscopy Core (RRID:SCR_019060), is supported, in part, by funding from the NIH-NINDS Neuroscience Center Support Grant P30 NS045892 and the NIH-NICHD Intellectual and Developmental Disabilities Research Center Support Grant P50 HD103573.

## Author contributions

J.L.S., D.F., K.H.T, S.Y., E.W. and M.G. conceived the experiments. M.G., E.W., and S.Y. generated all replicate organoid differentiations in this study. A.B. assisted in feeding and imaging the organoids. N.P. generated day 56 organoids. M.G. performed scRNA-seq library preparation. M.G. and M.L. analyzed scRNAseq data with N.M.’s assistance. E.W. performed flow cytometry experiments and analysis. S.Y. performed qPCR experiments and analysis. S.A., M.S., L.D., M.Y., and K.S. visually classified the organoids. T.F. supervised organoid ranking and area measurements, and annotated 3D organoid images. M.I.L. provided advice on statistical analysis. K.H.T, D.F., and J.L.S. supervised the work. M.G., E.W., S.Y., K.H.T, D.F., and J.L.S. wrote the manuscript. All authors read and approved the final manuscript.

---

**Supplementary Figure 1.**
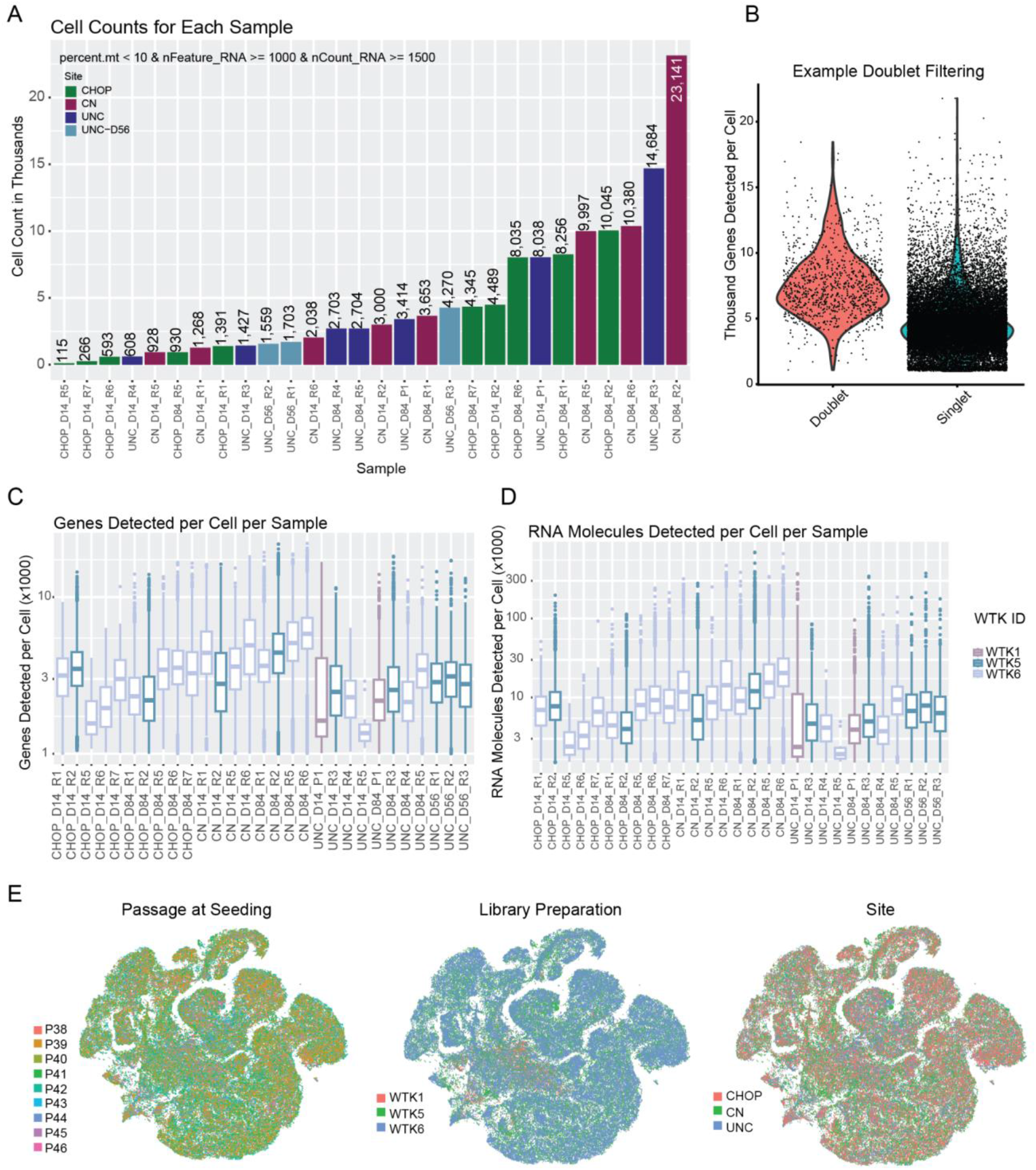
Single Cell RNA Sequencing Quality Control. A) Final cell counts for each sample in scRNAseq after all quality control. One day 14 sample (UNC R5) was removed from the experiment because less than 100 cells remained following quality control. Fewer cells were loaded into the library preparation at day 14 due to smaller organoid size and likely reduced heterogeneity, so the fewer cells observed at this time point are expected. B) DoubletFinder finds a higher number of genes detected per cell in predicted doublets as compared to predicted singlets. C) Average gene detected per cell and D) average RNA molecules detected per cell per sample, colored by Whole Transcriptome Kit (WTK) ID, which marks the library preparation batch. E) Colored tSNEs by technical batches and site.

**Supplemental Figure 2.**
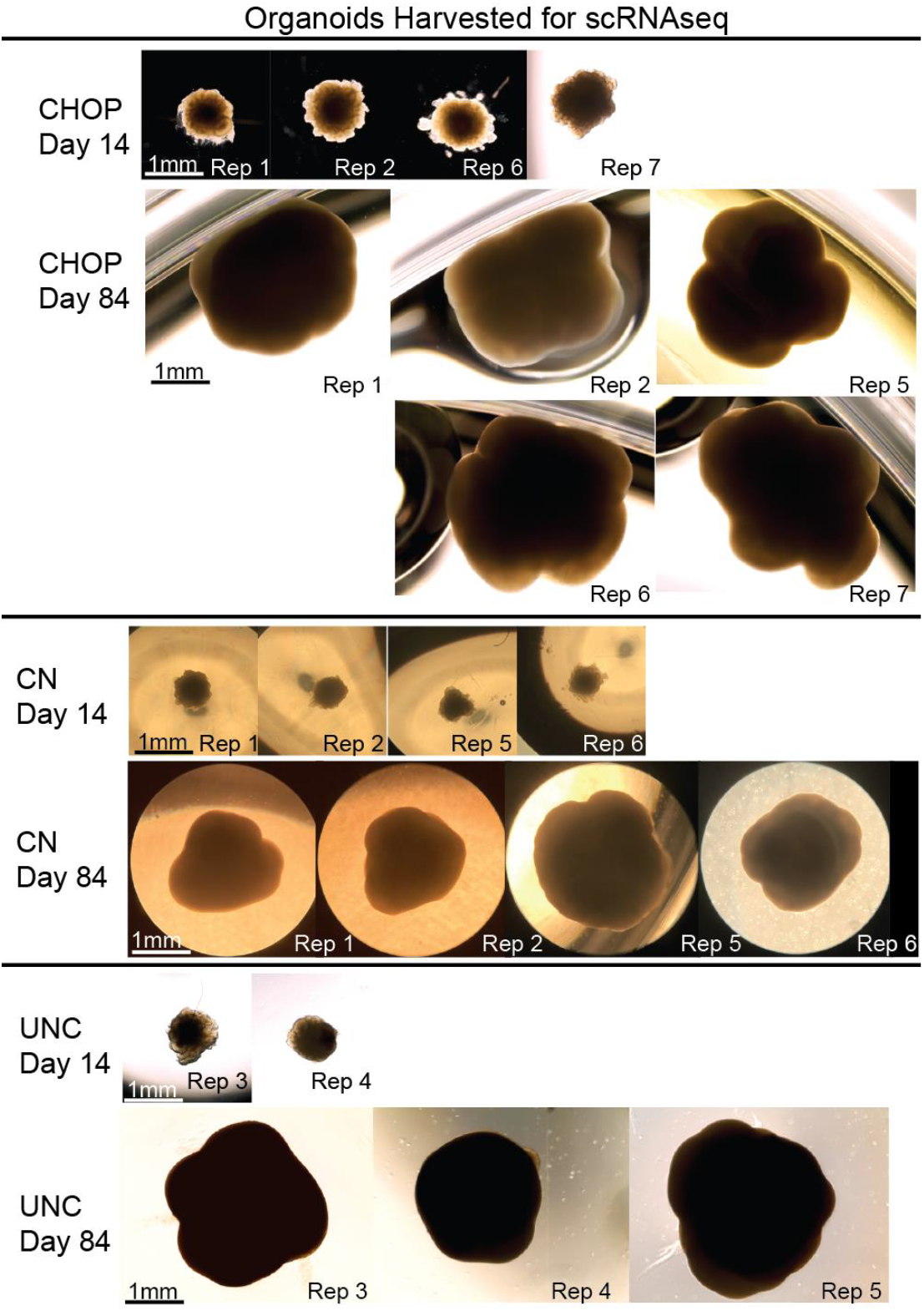
Images of Organoids Used for scRNAseq. CN size measurements are estimated due to imaging warping, and the sizes of these organoids are not directly comparable with the other two sites. UNC Rep 5 is not shown because scRNAseq samples did not pass quality control. UNC pilot 1 did not have phase contrast images collected, so is also not shown.

**Supplemental Figure 3.**
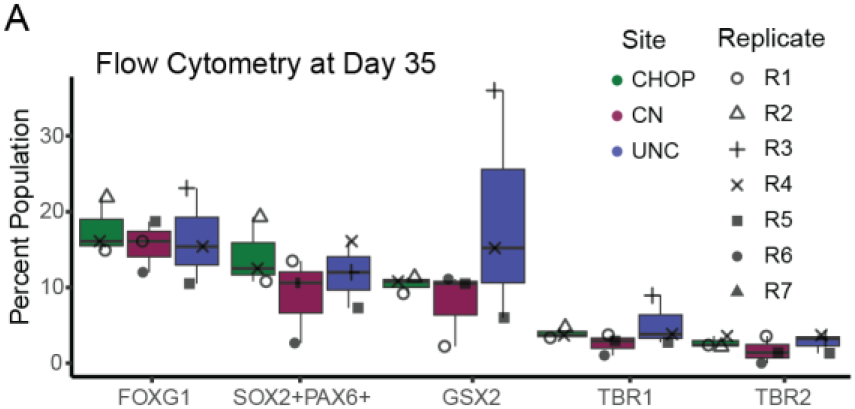
Percent population expressing cortical and non-cortical cell type markers at day 35. FOXG1, SOX2+/PAX6+ and TBR2 indicate cortical progenitors. GSX2 is a marker for ventral forebrain progenitors <u>(Corbin et al. 2000)</u>. TBR1 is a marker for early born cortical neurons. No significant differences across sites were detected.

**Supplementary Figure 4.**
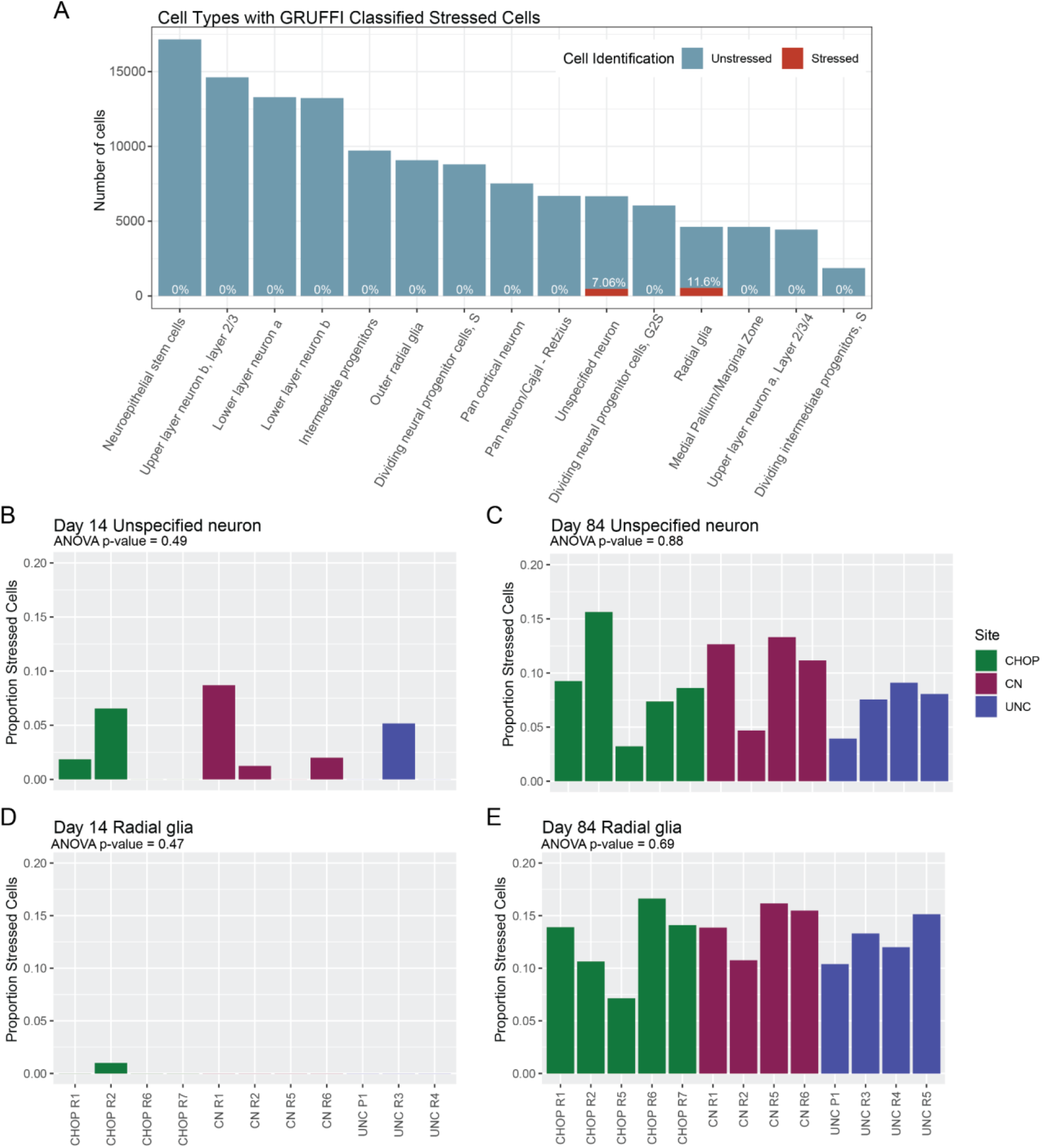
Gruffi Identified Cells under Metabolic Stress. A) Percent stressed cells across cell types. B) Proportion of stressed cells in each scRNAseq experiment in B) day 14 unspecified neurons C) day 84 unspecified neurons, D) day 14 radial glia and E) day 84 radial glia. No significant differences were detected across sites.

**Supplemental figure 5.**
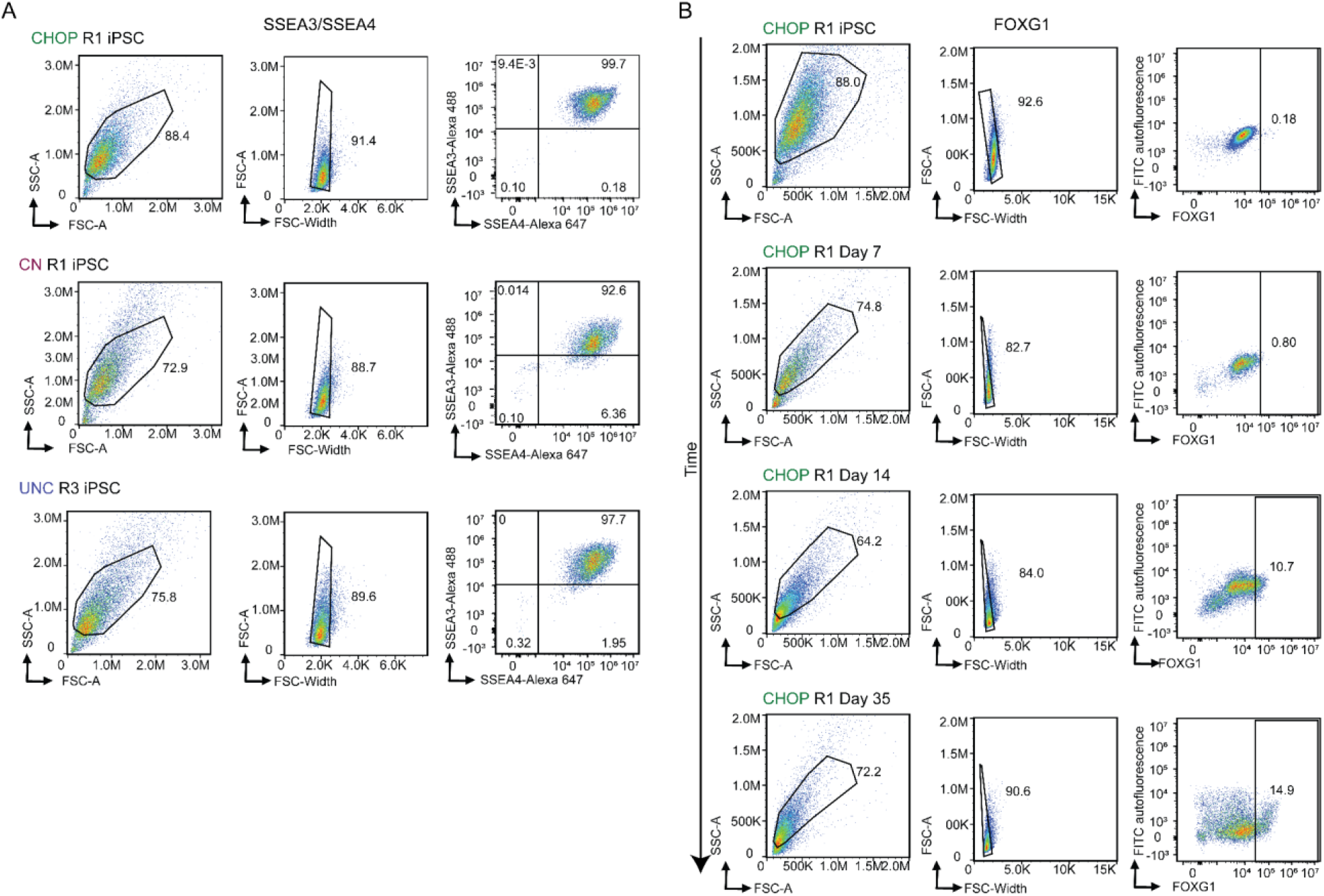
Example Flow Cytometry Gating. A) Gating for FOXG1 labeled cells in one sample across time. Gating was determined by day −1 immunoreactivity to FOXG1, which was negative. B) Gating for SSEA3/SSEA4 labeled cells from iPSC across all sites, which was determined relative to a no antibody control.

**Supplemental Figure 6.**
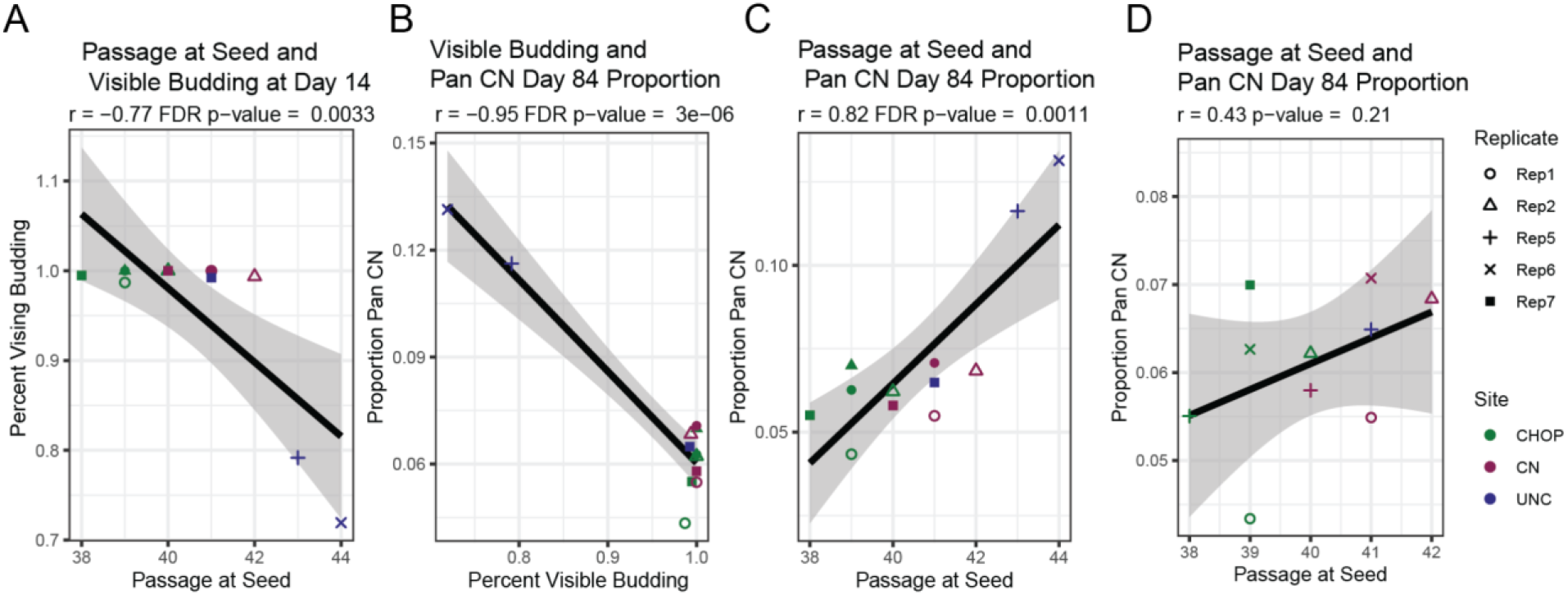
Correlates to day 84 cell type proportions. A) Pearson’s correlation between the two variables that correlated to the proportion of pan cortical neurons at day 84. Correlation between the proportion of pan cortical neurons at day 84 and B) percent visible budding or C) passage at seed. D) Correlation between passage at seed and the proportion of pan cortical neurons at day 84 when UNC R3 and UNC R4 outliers are removed.

