## Supplemental material for "Cross-site reproducibility of human cortical organoids reveals consistent cell type composition and architecture"

**Supplemental Table 1. Experiments Included in Each Analysis.** Number in the parentheses indicates the number of hCOs used, if applicable. X indicates if the experiment was included in each assay otherwise.

| Replicate Differentiation | Notes | Flow iPSC | qPCR iPSC | Imaging Day 14 | scRNAseq Day 14 | Flow Day 35 | scRNAseq Day 84 | Imaging Large hCO Day 84 | Imaging Small hCO Day 84 |
| --- | --- | --- | --- | --- | --- | --- | --- | --- | --- |
| UNC P1 | Completed |  |  | 3 | 1 |  | 1 |  |  |
| UNC R1 | Mostly cysts day 35 |  |  |  |  |  |  |  |  |
| UNC R2 | Poor seeding |  |  |  |  |  |  |  |  |
| UNC R3 | Completed | x | x | 3 | 1 | 2 | 1 | 3 | 3 |
| UNC R4 | Completed | x | x | 3 | 1 | 2 | 1 |  | 3 |
| UNC R5 | Completed | x | x | 1 |  | 2 | 1 | 2 |  |
| UNC R6 | Mostly cysts day 35 |  |  |  |  |  |  |  |  |
| UNC R7 | Mostly cysts day 35 |  |  |  |  |  |  |  |  |
| CHOP R1 | Completed | x | x | 3 | 1 | 2 | 1 |  |  |
| CHOP R2 | Completed | x | x | 3 | 1 | 2 | 1 |  | 3 |
| CHOP R3 | Poor seeding |  |  |  |  |  |  |  |  |
| CHOP R4 | Dying D56+ | x | x | 4 |  | 2 |  |  |  |
| CHOP R5 | Completed | x | x |  |  |  | 1 | 2 | 2 |
| CHOP R6 | Completed | x | x |  | 1 |  | 1 | 1 |  |
| CHOP R7 | Completed | x | x |  | 1 |  | 1 |  |  |
| CN R1 | Completed | x | x | 2 | 1 | 2 | 1 |  |  |
| CN R2 | Completed | x | x | 3 | 1 |  | 1 |  | 3 |
| CN R3 | Contamination day 70 |  | x |  |  |  |  |  |  |
| CN R4 | Contamination day 70 |  | x | 2 |  |  |  |  |  |
| CN R5 | Completed | x | x | 1 | 1 | 2 | 1 | 2 |  |
| CN R6 | Completed | x | x | 1 | 1 | 2 | 1 | 2 | 3 |
| CN R7 | Contamination day 84 | x | x |  |  |  |  |  |  |

**Supplemental Table 2. Cell Type Proportions Across Site at Day 14.** Baseline proportion is the geometric mean across all samples.

| Cell Type | Baseline Proportion | CHOP Mean Proportion | CN Mean Proportion | UNC Mean Proportion | F statistic | p-value | FDR adjusted p-value |
| --- | --- | --- | --- | --- | --- | --- | --- |
| Neuroepithelial stem cells | 0.524351 | 0.509456 | 0.461311 | 0.513454 | 0.222612 | 0.803204 | 0.919217 |
| Dividing neural progenitor cells, S | 0.01599 | 0.004674 | 0.005228 | 0.018194 | 2.595466 | 0.112027 | 0.420099 |
| Dividing neural progenitor cells, G2 | 0.008275 | 0.003856 | 0.003905 | 0.011181 | 1.312114 | 0.302157 | 0.735733 |
| Medial Pallium/Marginal Zone | 0.030515 | 0.027257 | 0.032796 | 0.045625 | 0.72294 | 0.503567 | 0.75535 |
| Dividing intermediate progenitors, S | 0.015731 | 0.014251 | 0.014262 | 0.013051 | 0.114712 | 0.892452 | 0.919217 |
| Intermediate progenitors | 0.003017 | 0.002246 | 0.004381 | 0.001808 | 0.986119 | 0.398883 | 0.735733 |
| Radial glia | 0.168779 | 0.211618 | 0.221383 | 0.160019 | 0.867378 | 0.44144 | 0.735733 |
| Outer radial glia | 0.005258 | 0.002219 | 0.003109 | 0.007658 | 1.078077 | 0.368388 | 0.735733 |
| Lower layer neuron a | 0.057797 | 0.060844 | 0.065382 | 0.051786 | 0.183683 | 0.834173 | 0.919217 |
| Lower layer neuron b | 0.024955 | 0.032326 | 0.030436 | 0.030516 | 0.084772 | 0.919217 | 0.919217 |
| Upper layer neuron a, Layer 2/3/4 | 0.072623 | 0.088935 | 0.076802 | 0.065337 | 0.423934 | 0.662593 | 0.903535 |
| Upper layer neuron b, layer 2/3 | 0.023188 | 0.020867 | 0.040569 | 0.01453 | 2.730464 | 0.101751 | 0.420099 |
| Pan cortical neuron | 0.043832 | 0.020042 | 0.038346 | 0.058746 | 3.449677 | 0.062324 | 0.420099 |
| Pan neuron/Cajal - Retzius | 0.005086 | 9.62E-04 | 0.001787 | 0.007148 | 5.476333 | 0.01823 | 0.273445 |
| Unspecified neuron | 6.03E-04 | 4.45E-04 | 3.02E-04 | 9.47E-04 | 1.318281 | 0.300607 | 0.735733 |

Supplemental Table 3. Cell Type Proportions Across Site at Day 84. Baseline proportion is the geometric mean across all samples.

| Cell Type | Baseline Proportion | CHOP Mean Proportion | CN Mean Proportion | UNC Mean Proportion | F statistic | p-value | FDR adjusted p-value |
| --- | --- | --- | --- | --- | --- | --- | --- |
| Neuroepithelial stem cells | 0.049087 | 0.042896 | 0.052552 | 0.044627 | 0.545451 | 0.591776 | 0.73972 |
| Dividing neural progenitor cells, S | 0.135716 | 0.144869 | 0.132356 | 0.137808 | 0.19155 | 0.827874 | 0.955239 |
| Dividing neural progenitor cells, G2 | 0.123917 | 0.126196 | 0.131555 | 0.117632 | 0.628828 | 0.548065 | 0.73972 |
| Medial Pallium/Marginal Zone | 0.120752 | 0.113855 | 0.134178 | 0.103815 | 1.971152 | 0.177217 | 0.443042 |
| Dividing intermediate progenitors, S | 0.087579 | 0.102227 | 0.084289 | 0.083515 | 1.08255 | 0.366275 | 0.610458 |
| Intermediate progenitors | 0.0827 | 0.081275 | 0.092048 | 0.078186 | 0.835734 | 0.4547 | 0.68205 |
| Radial glia | 0.042881 | 0.044182 | 0.048275 | 0.025052 | 8.201397 | 0.004626 | 0.060919 |
| Outer radial glia | 0.072972 | 0.058644 | 0.062994 | 0.09927 | 6.131636 | 0.01268 | 0.0634 |
| Lower layer neuron a | 0.052782 | 0.050621 | 0.051075 | 0.051338 | 0.017467 | 0.982707 | 0.982707 |
| Lower layer neuron b | 0.055599 | 0.047656 | 0.048669 | 0.076248 | 1.761808 | 0.210082 | 0.450176 |
| Upper layer neuron a, Layer 2/3/4 | 0.040115 | 0.043807 | 0.042423 | 0.031818 | 2.811542 | 0.095166 | 0.285497 |
| Upper layer neuron b, layer 2/3 | 0.039768 | 0.041643 | 0.033774 | 0.050226 | 2.931896 | 0.08745 | 0.285497 |
| Pan cortical neuron | 0.034766 | 0.038648 | 0.025539 | 0.043923 | 1.175797 | 0.339124 | 0.610458 |
| Pan neuron/Cajal - Retzius | 0.043718 | 0.041988 | 0.042608 | 0.044157 | 0.038933 | 0.961926 | 0.982707 |
| Unspecified neuron | 0.017648 | 0.021494 | 0.017667 | 0.012384 | 7.008118 | 0.008123 | 0.060919 |

Resources Table 1. qPCR Primers.

| Resource | Company | Citation |
| --- | --- | --- |
| EIF4A2 fw 5’-TGGTGTCATCGAGAGCAACTG-3’ rv 5’-GGCTTCTCAAAACCGTAAGCA-3’ | Integrated DNA Technologies | (Watanabe et al. 2022) |
| Nanog fw 5’-TTTGTGGGCCTGAAGAAAACT-3’ rv 5’-AGGGCTGTCCTGAATAAGCAG-3’ | Integrated DNA Technologies | (Watanabe et al. 2022) |
| OCT4 fw 5’-GGAGAAGCTGGAGCAAAAC-3’ rv 5’-ACCTTCCCAAATAGAACCCC-3’ | Integrated DNA Technologies | (Watanabe et al. 2022) |
| OTX2 fw 5’-AGAGGACGACGTTCACTCG-3’ rv 5’-TCGGGCAAGTTGATTTTCAGT-3’ | Integrated DNA Technologies | (Collier et al. 2017) |
| ZIC2 fw 5’-GATGTGCGACAAGTCCTACAC-3’ rv 5’-TGGACGACTCATAGCCGGA-3’ | Integrated DNA Technologies | (Collier et al. 2017) |
| DUSP6 fw 5’-ACAAGCAAATCCCCATCTCG-3’ rv 5’-CAGCCAAGCAATGTACCAAG-3’ | Integrated DNA Technologies | (Alves et al. 2015) |
| KLF4 fw 5’-GCTGCCGAGGACCTTCTG-3’ rv 5’-GCGAACGTGGAGAAAGATGG-3’ | Integrated DNA Technologies | (Watanabe et al. 2022) |
| TFAP2C fw 5’-CTGTTGCTGCACGATCAGACA-3’ rv 5’-CTCAGTGGGGTTCATTACGGC-3’ | Integrated DNA Technologies | (Watanabe et al. 2022) |
| TFCP2L1 fw 5’-ATACCAGCCGTCCTATGAAACC-3’ rv 5’-ACTGCGAGAACCTGTTGCG-3’ | Integrated DNA Technologies | (Watanabe et al. 2022) |

Resources Table 2. Flow Cytometry Antibodies.

| **Target** | **Company** | **Catalog #** | **RRID** | **Dilution** |
| --- | --- | --- | --- | --- |
| SSEA3 | Biolegend | 330306 | RRID:AB_1279440 | 1:50 |
| SSEA4 | Biolegend | 330408 | RRID:AB_1089200 | 1:50 |
| PODXL | ThermoFisher | 12-8873-42 | RRID:AB_10734225 | 1:50 |
| FOXG1 | Abcam | ab196868 | RRID:AB_2892604 | 1:300 |
| SOX2 | Cell Signaling Technology | 3579 | RRID:AB_2195767 | 1:300 |
| PAX6 | BD | 562249 | RRID:AB_11152956 | 1:20 |
| TBR2 (EOMES) | Cell Signaling Technology | 81493 | RRID:AB_2799974 | 1:1000 |
| TBR1 | Proteintech, purchased through ThermoFisher | 20932-1-AP | RRID:AB_10695502 | 1:1000 |
| GSX2 (Gsh2) | Millipore | ABN162 | RRID:AB_11203296 | 1:1000 |
| **Secondary antibodies** |  |  |  |  |
| goat-anti-rabbit Alexa647 | Jackson Immunoresearch | 111-605-144 | RRID:AB_2338078 | 1:500 |
| goat-anti-rabbit Alexa488 | Jackson Immunoresearch | 111-545-144 | RRID:AB_2338052 | 1:500 |

Resources Table 3. 3D Imaging of iDISCO Processed Samples Antibodies.

| **Target** | **Company** | **Catalog #** | **RRID** | **Dilution** |
| --- | --- | --- | --- | --- |
| PAX6 | Thermo Fisher Scientific | PA5-85374 | RRID:AB_2792516 | 1:100 |
| NCAD | BD Biosciences | 610921 | RRID:AB_398236 | 1:200 |
| CTIP2 | Abcam | ab18465 | RRID:AB_18465 | 1:200 |
| **Secondary antibodies** |  |  |  |  |
| To-PRO-3 | Thermo Fisher Scientific | T3605 |  | 1:750 |
| Alexa Fluor goat anti-rat-568 | Thermo Fisher Scientific | A1107 | RRID:AB_2338052 | 1:750 |
| Alexa Fluor goat anti-rabbit-790 | Thermo Fisher Scientific | A11369 | RRID:AB_2534142 | 1:750 |
| Alexa Fluor goat anti-rabbit-405 | Thermo Fisher Scientific | A31556 | RRID:AB_221605 | 1:750 |
| Alexa Fluor goat anti-mouse-488 | Thermo Fisher Scientific | A28175 | RRID:AB_2536161 | 1:750 |

**Supplemental Table for Figure 5. Organoid Rank Scales Provided for Training.** Images are not to scale and are from differentiations not included in this study.

Day 14 Organoid Morphology Scale for Budding

| hCOs with visible budding | 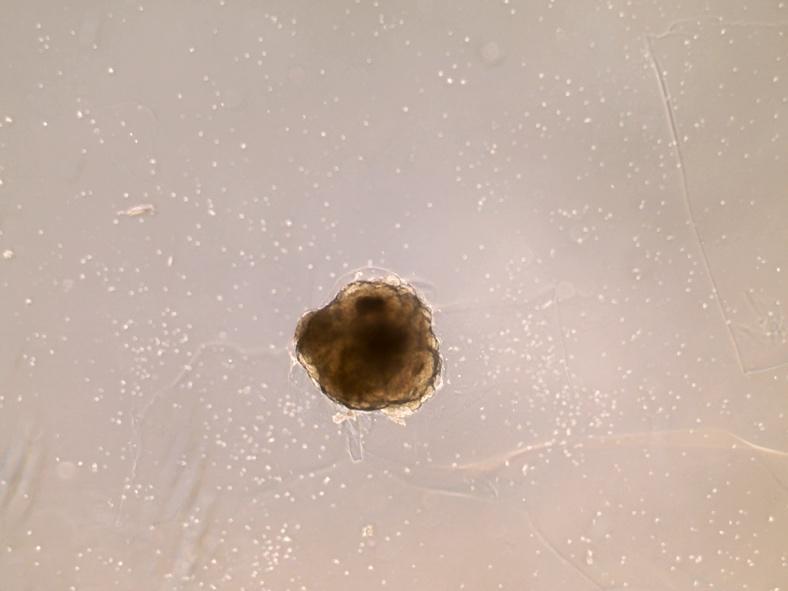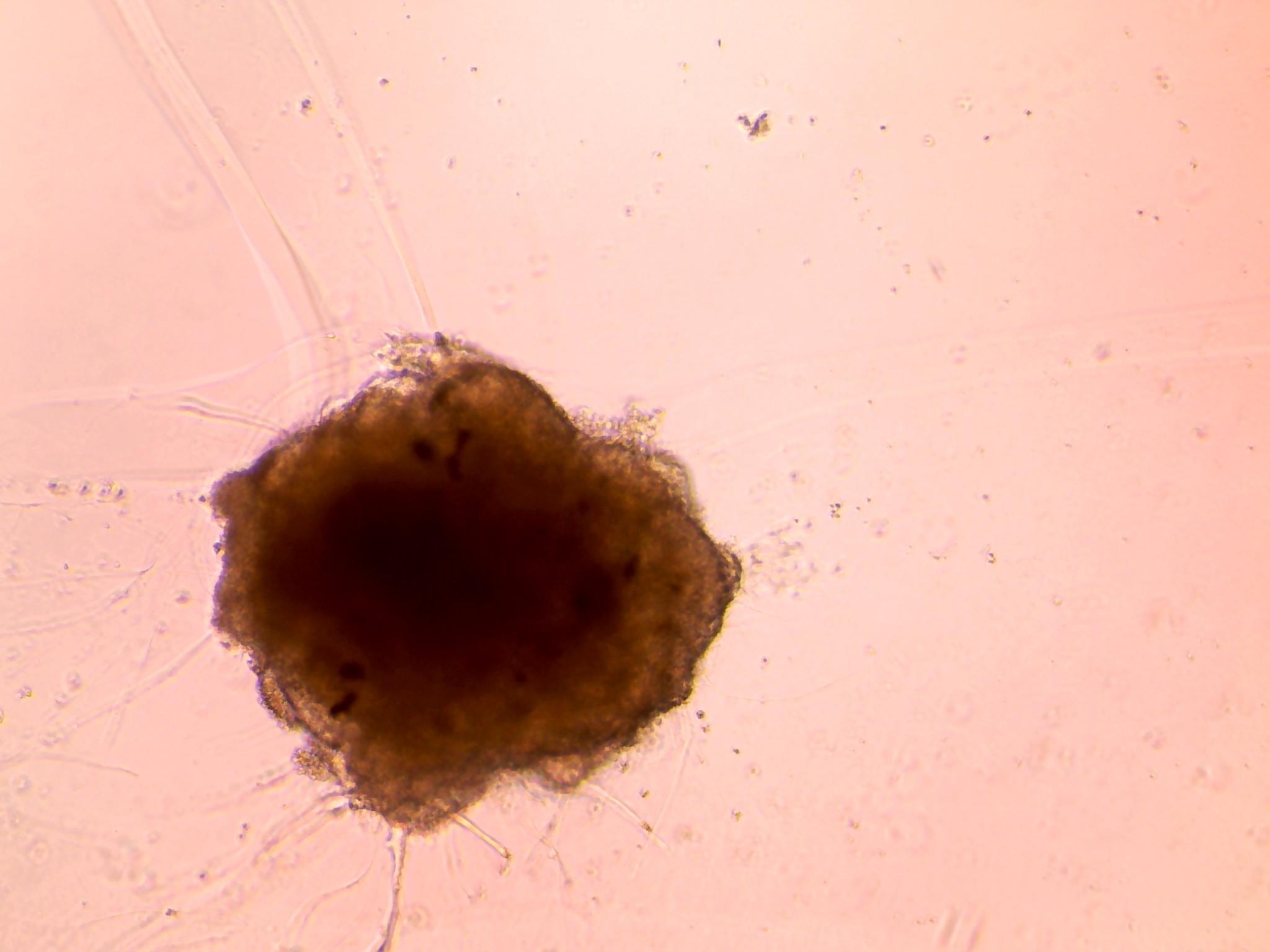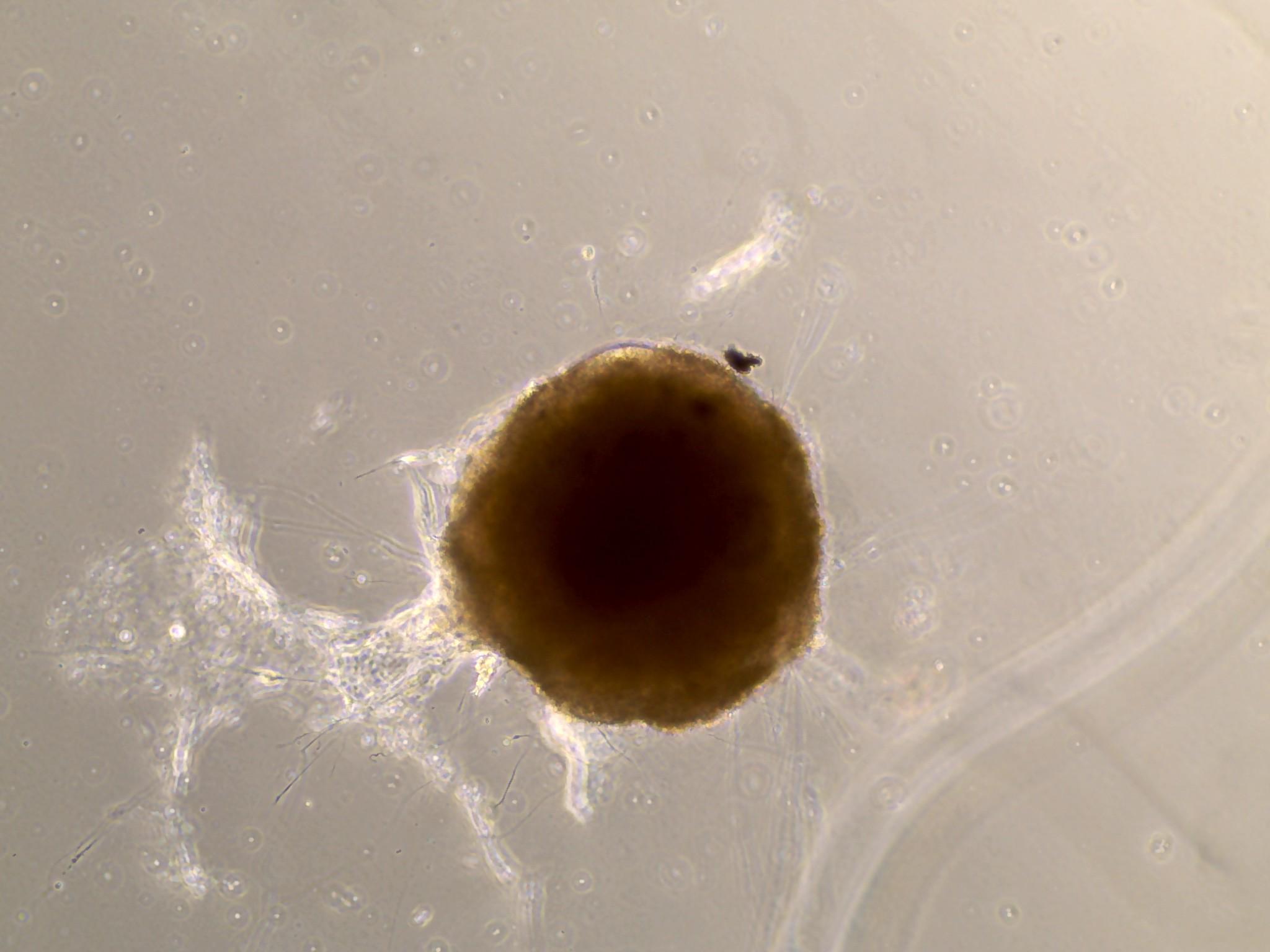 |
| --- | --- |
| hCOs without visible budding | 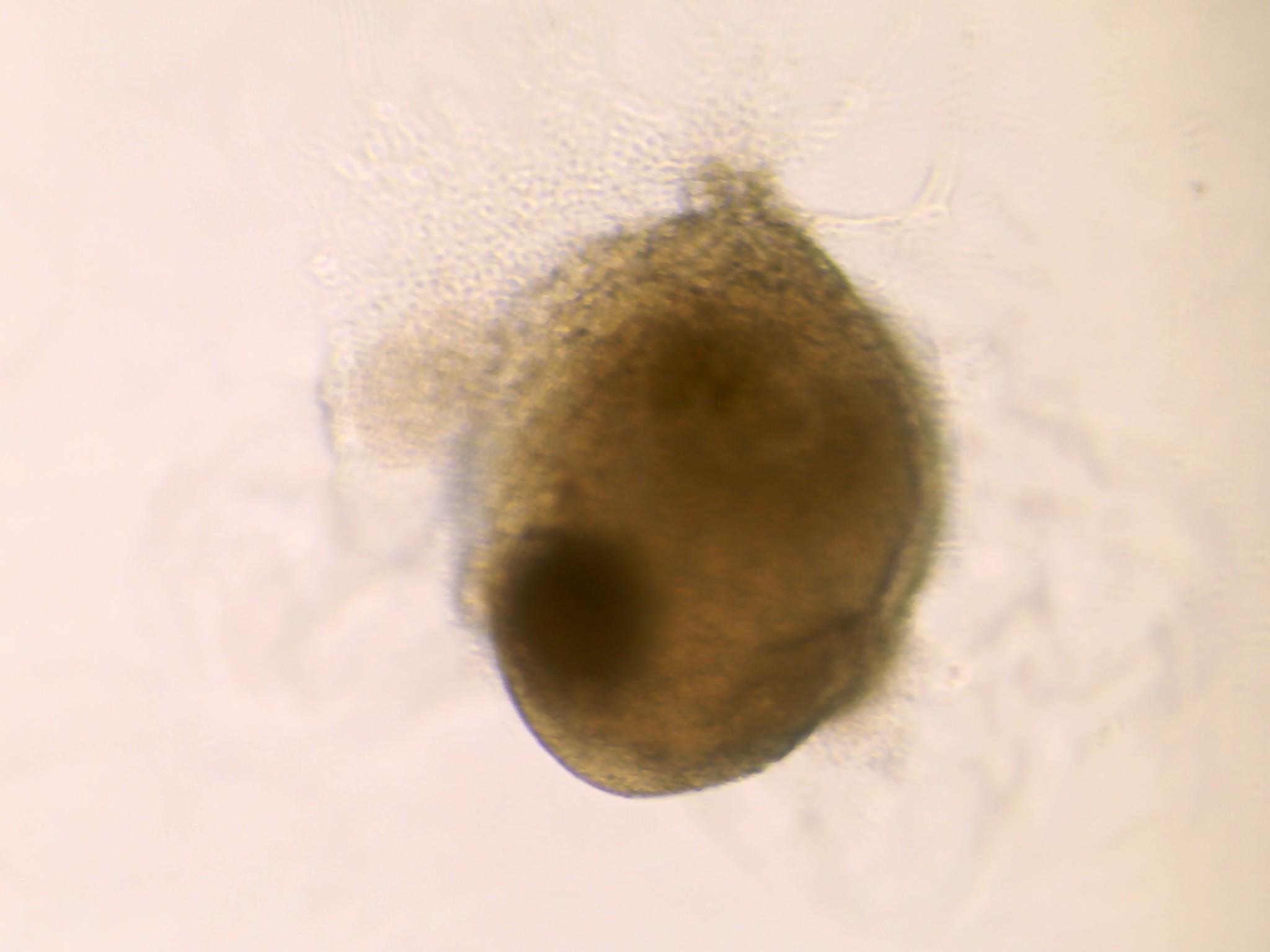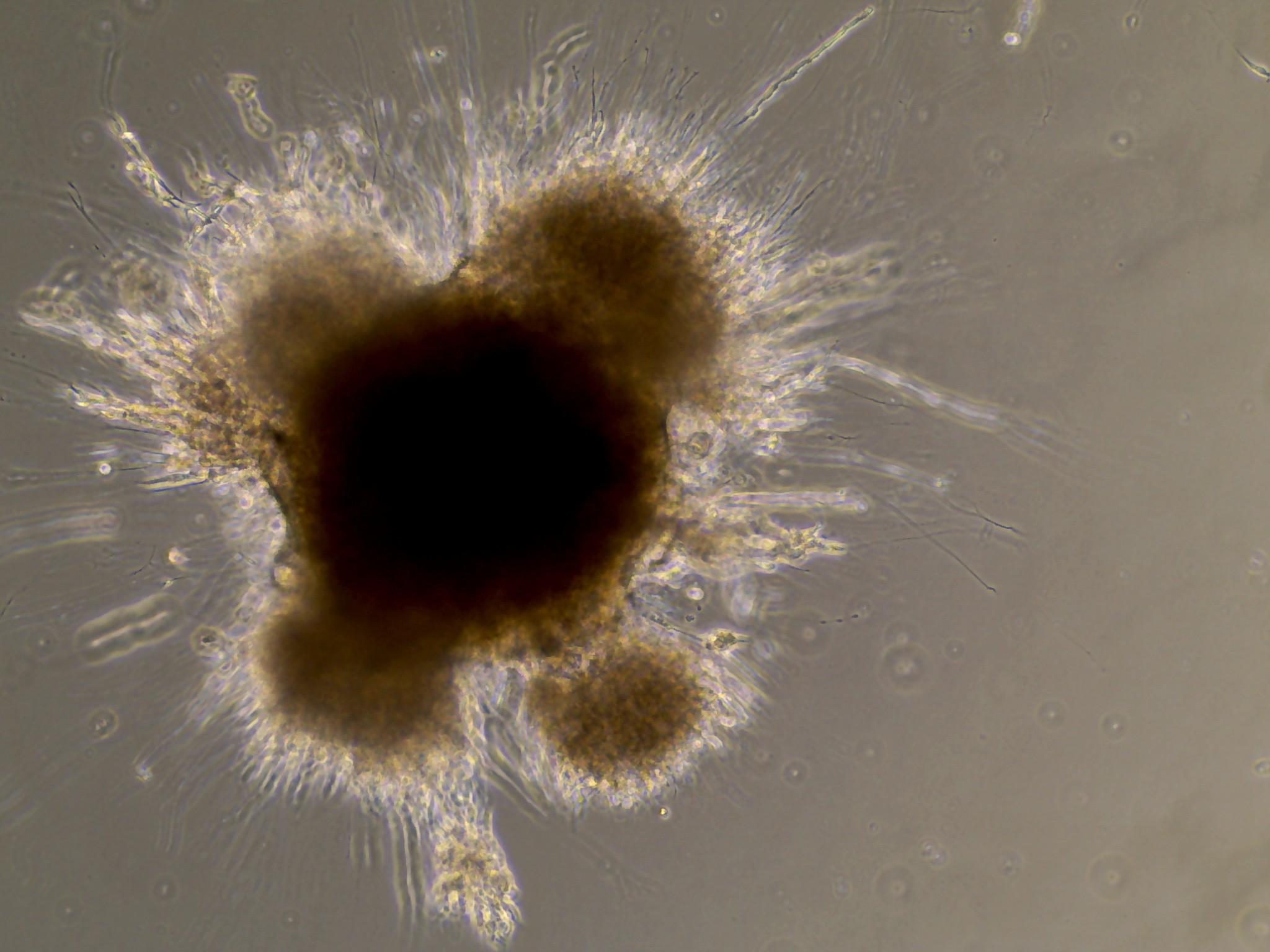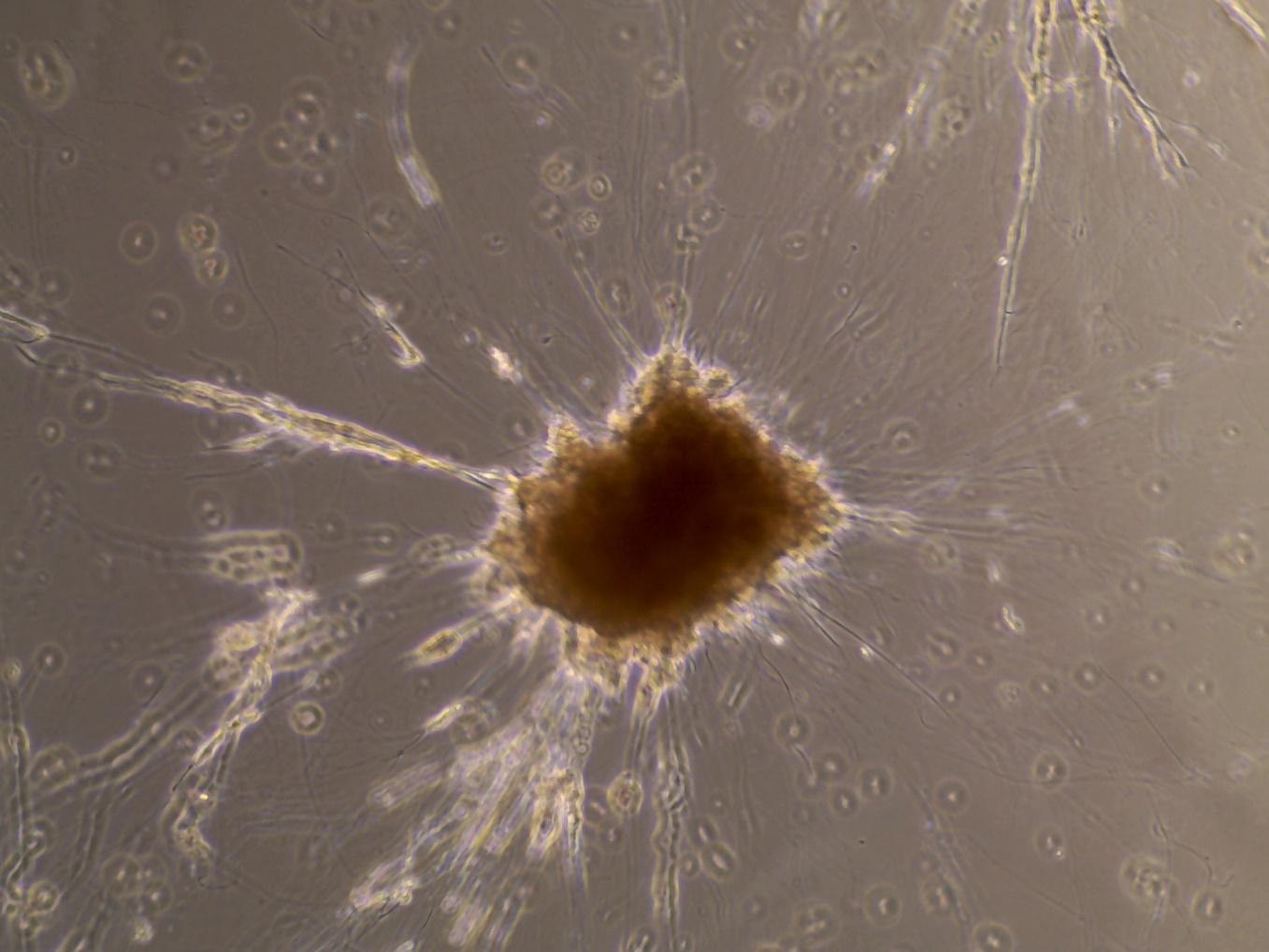 |

Day 14 Organoid Morphology Scale for Budding

| hCOs with no growth in matrigel | 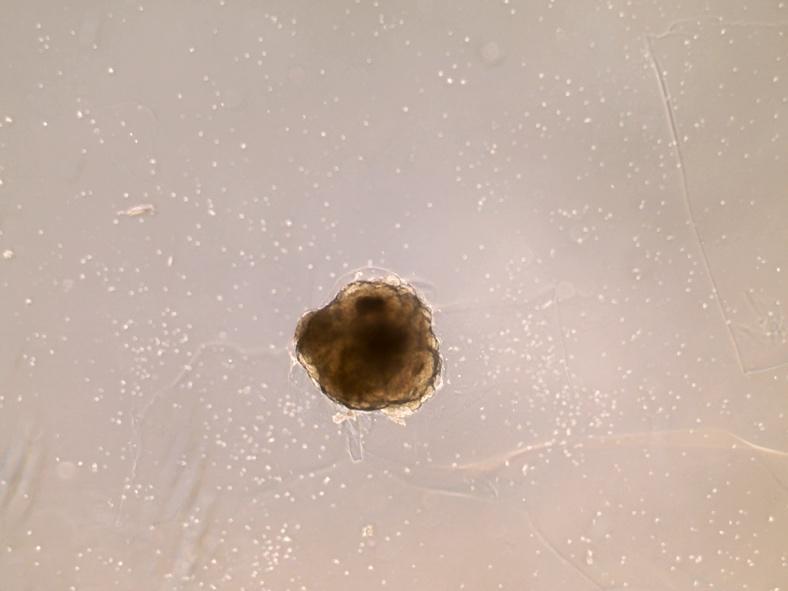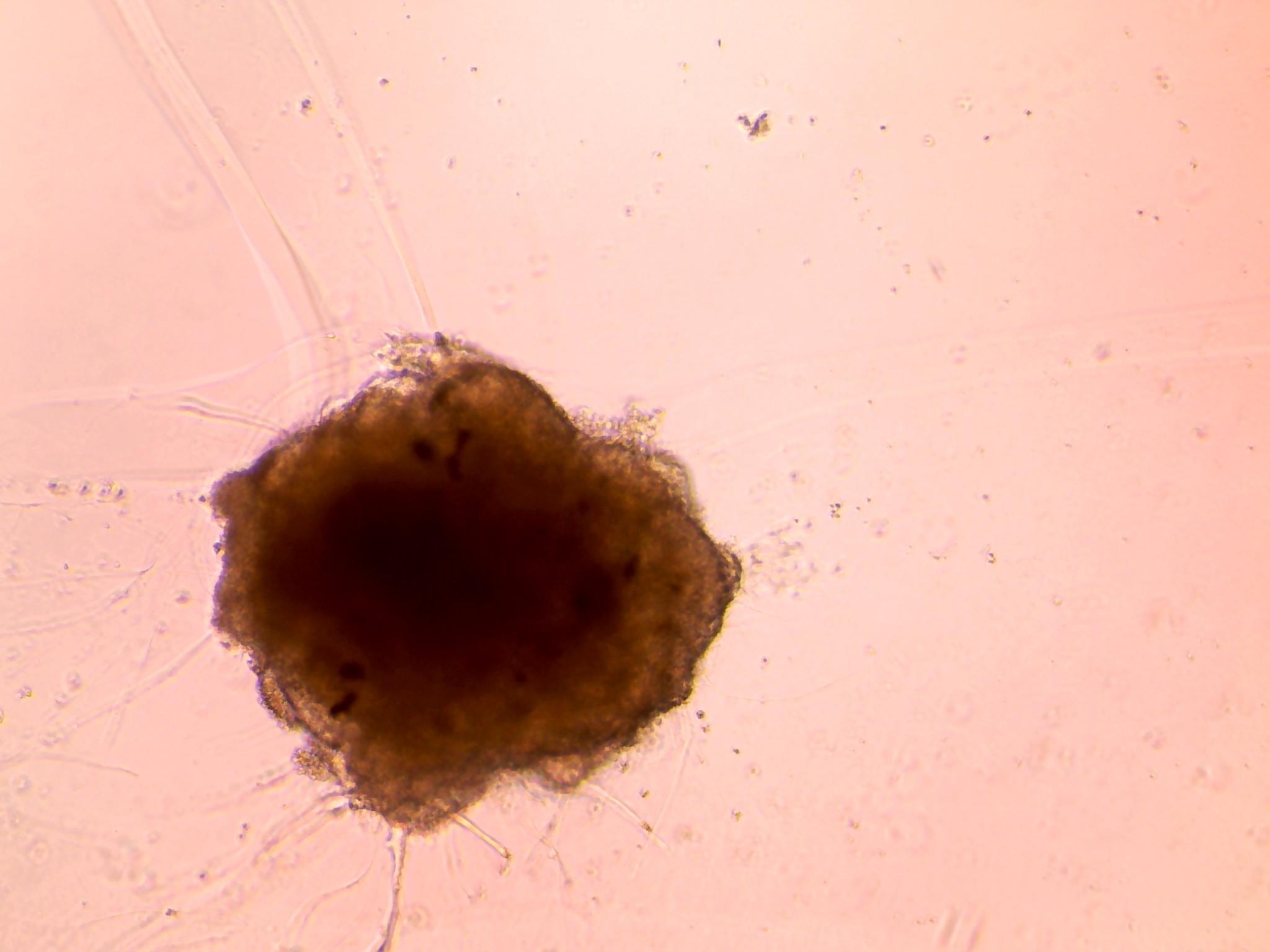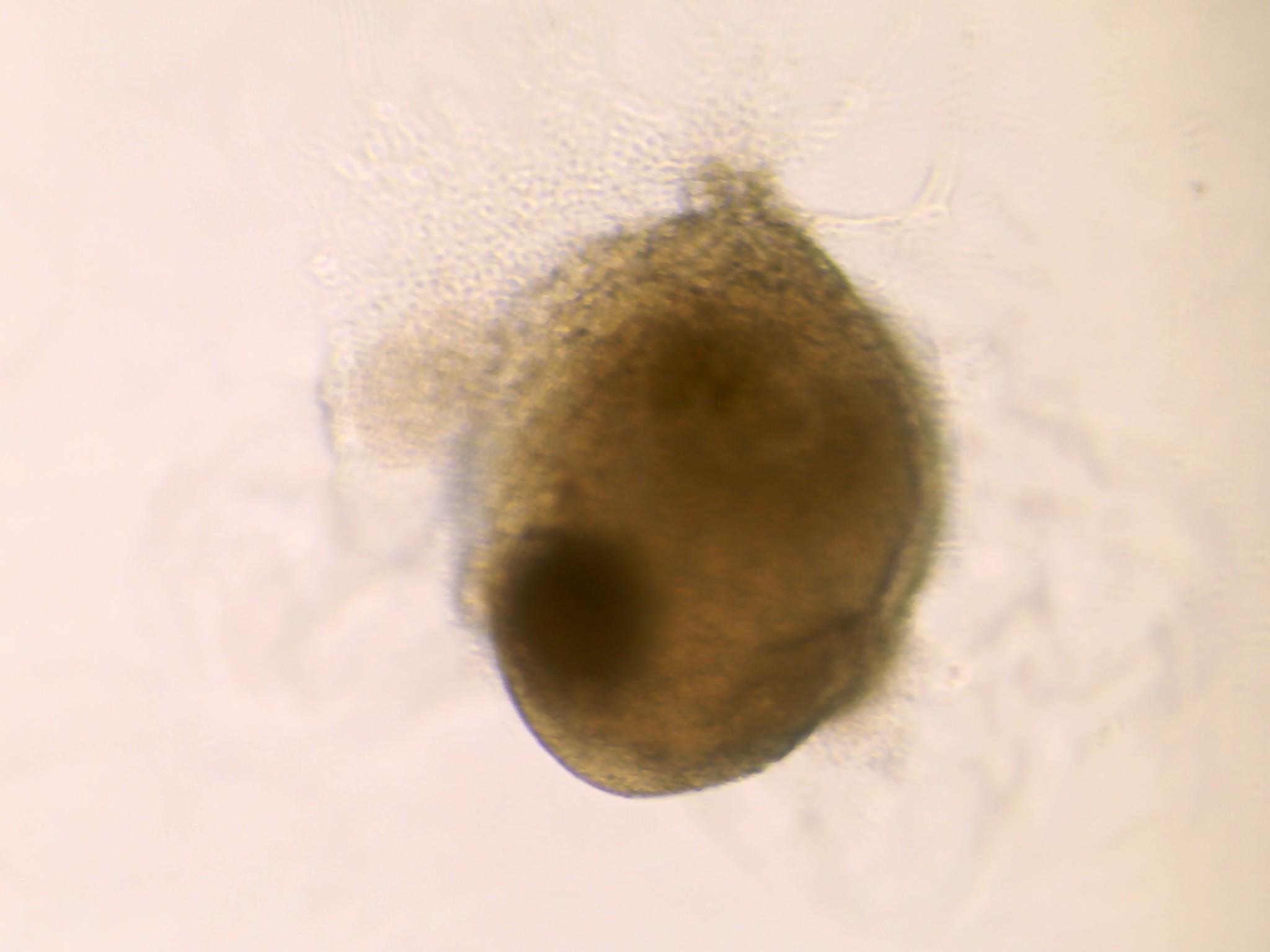 |
| --- | --- |
| hCOs with growth in matrigel | 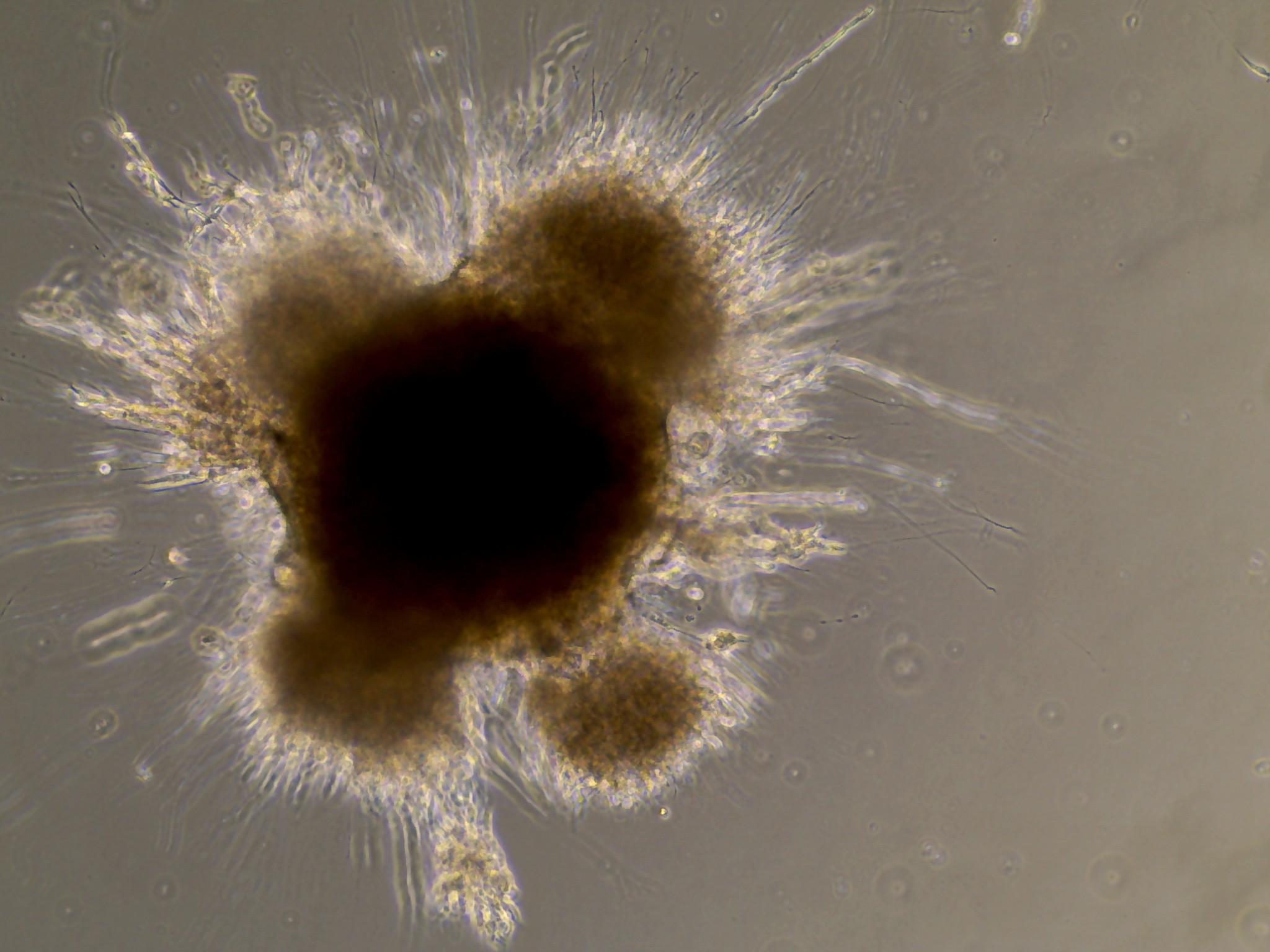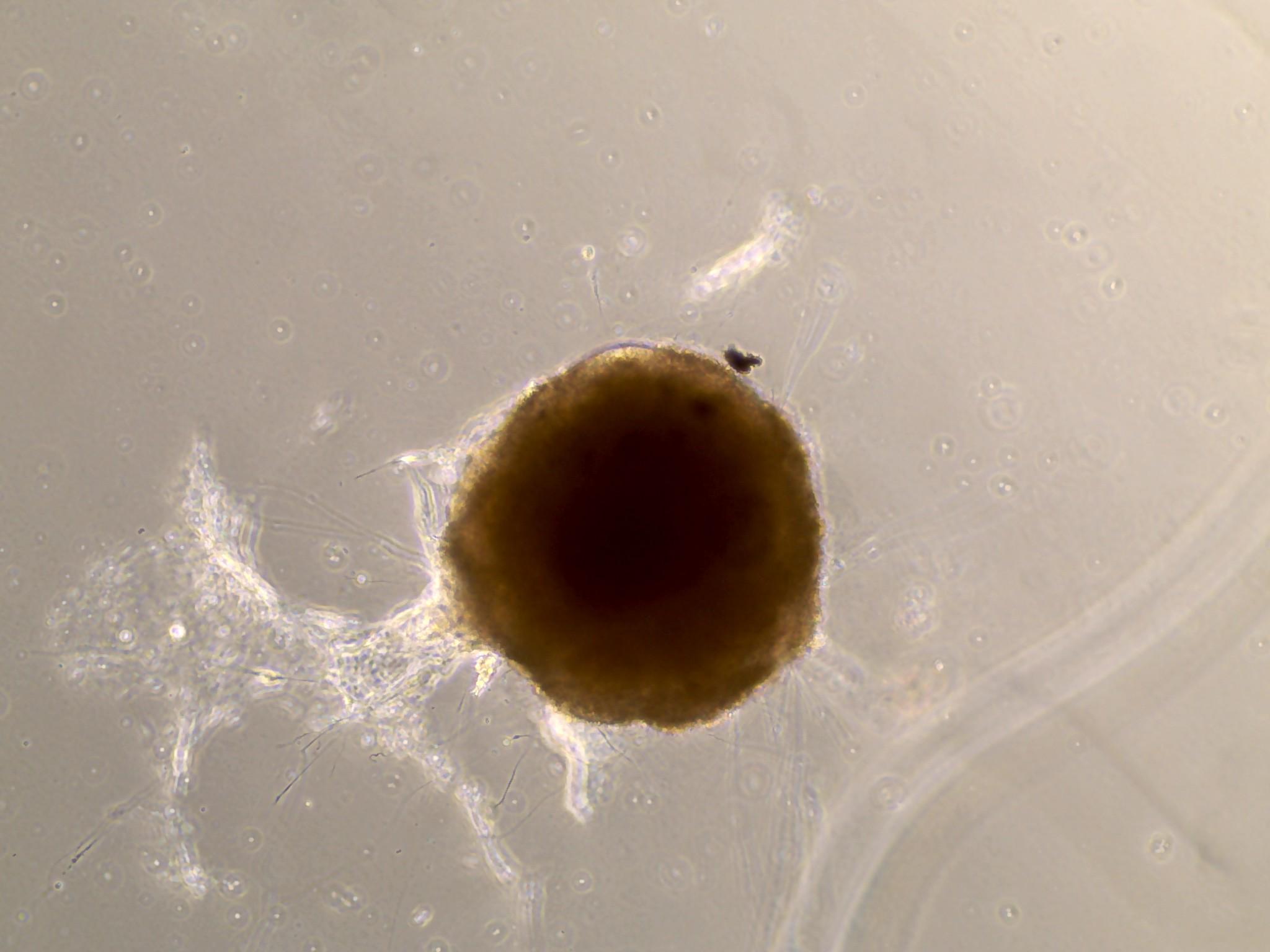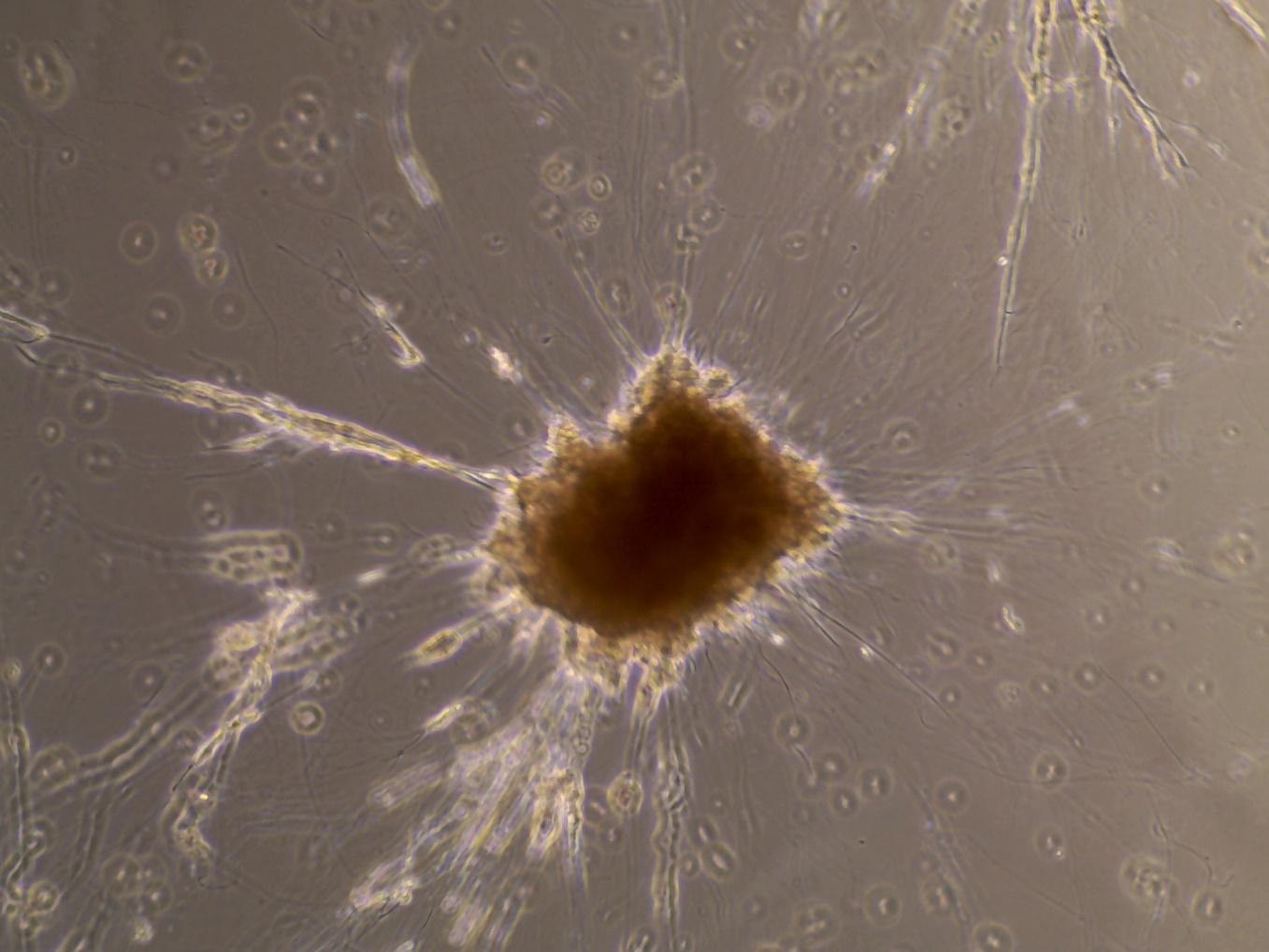 |

Day 35 Organoid Morphology Scale

| hCO with expected morphology | 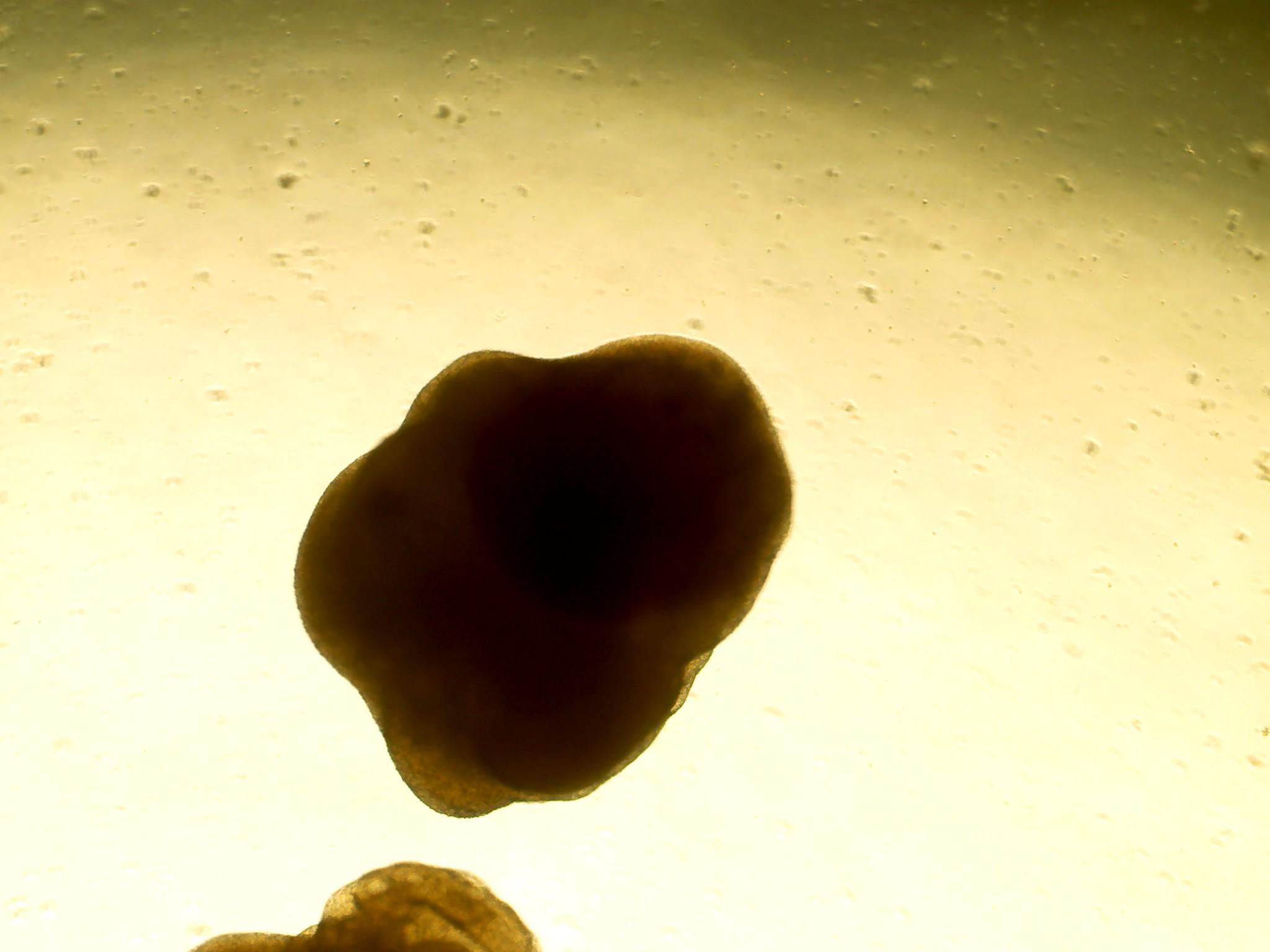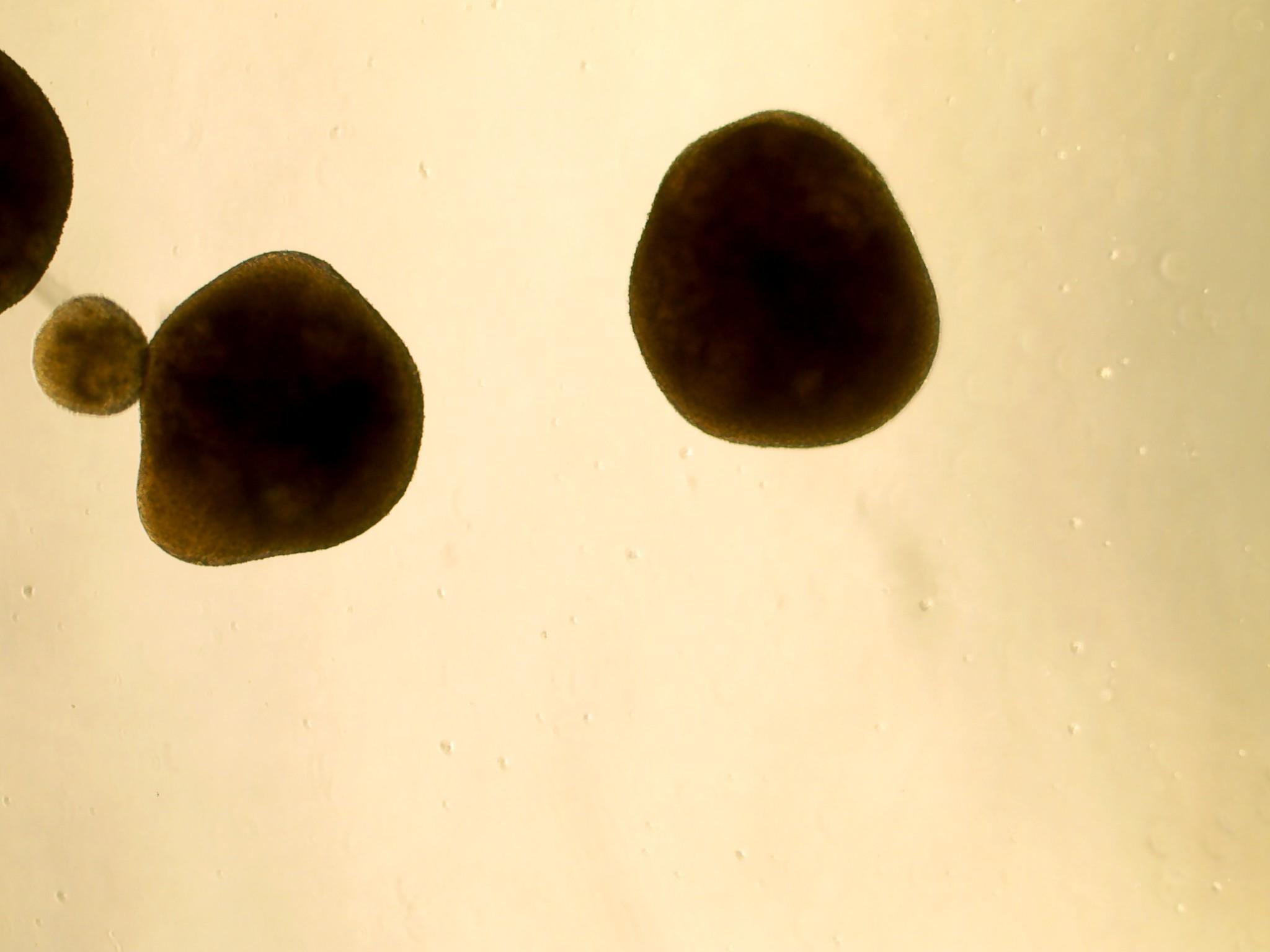 |
| --- | --- |
| hCO with unexpected morphology | 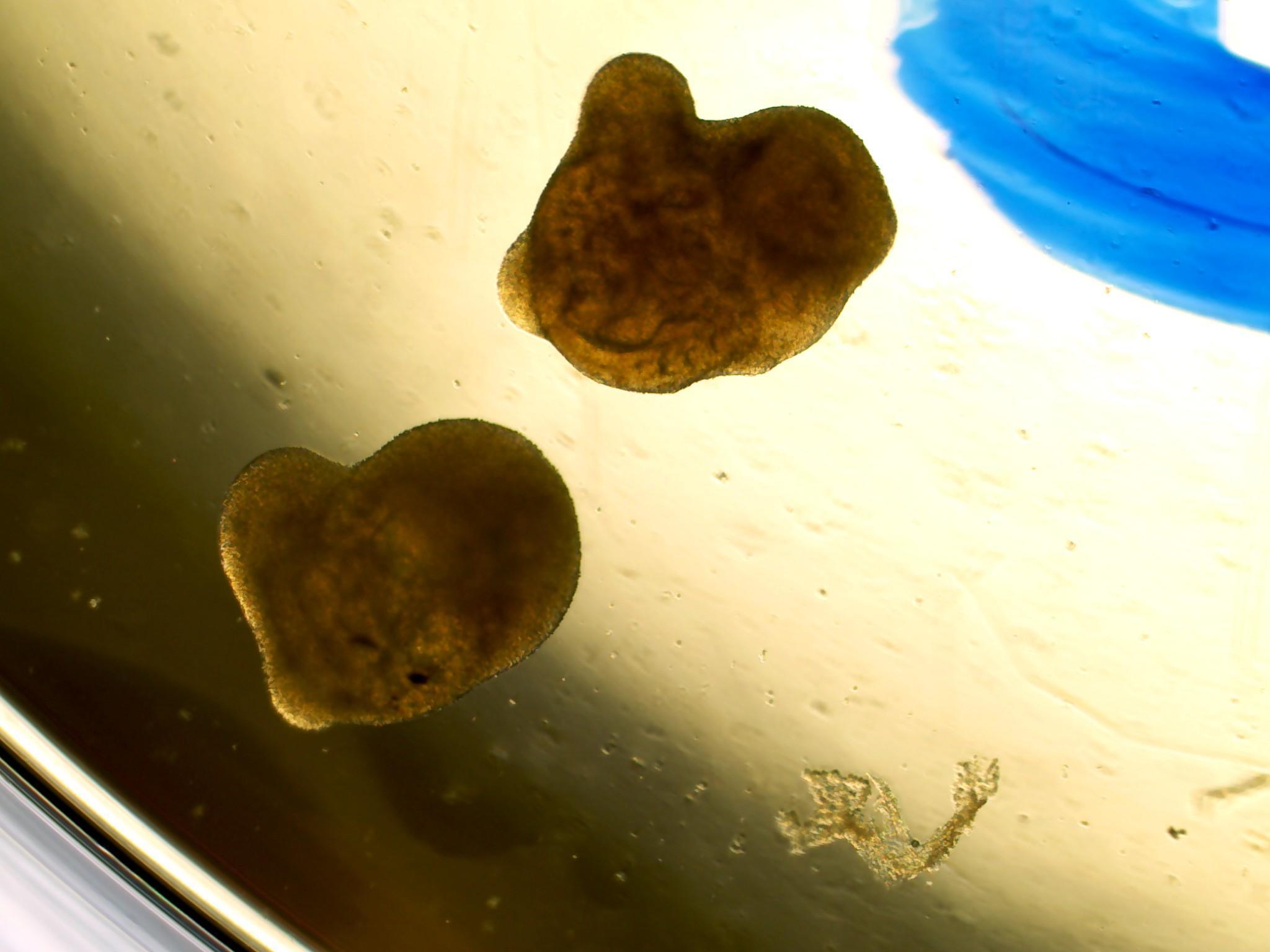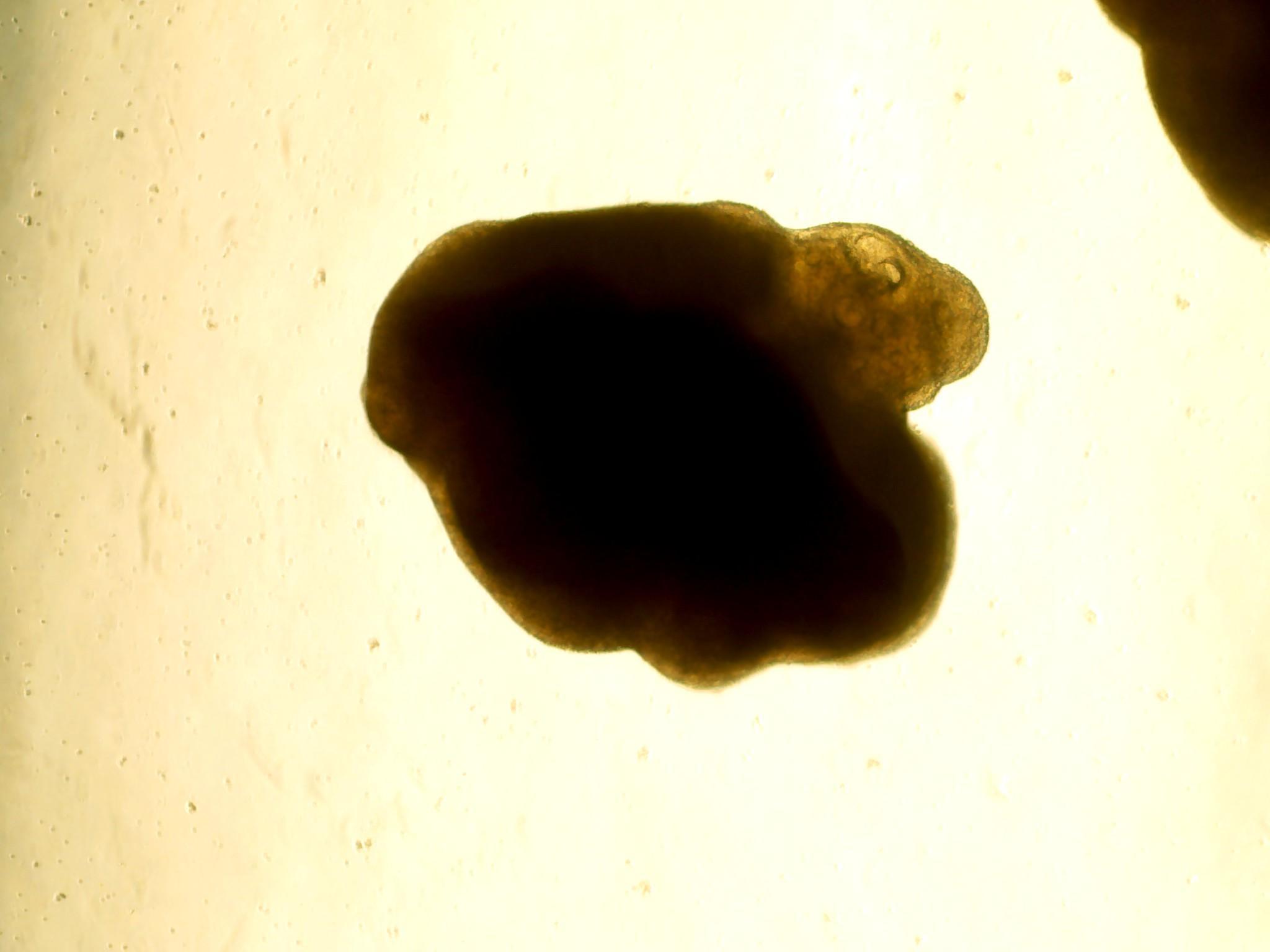 |

Day 56 Organoid Morphology Scale

| hCO with expected morphology | 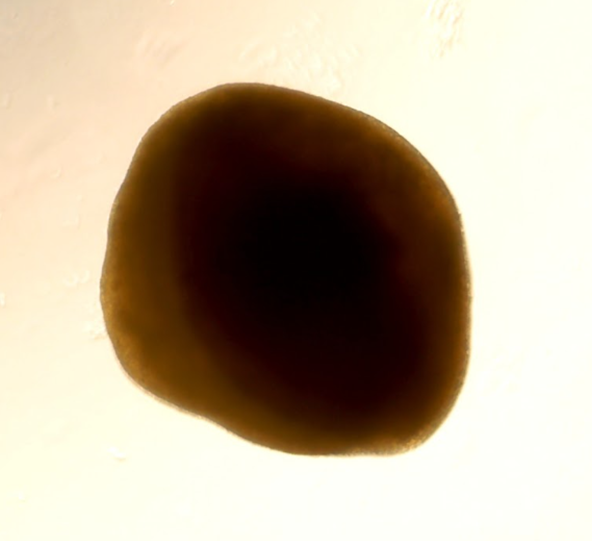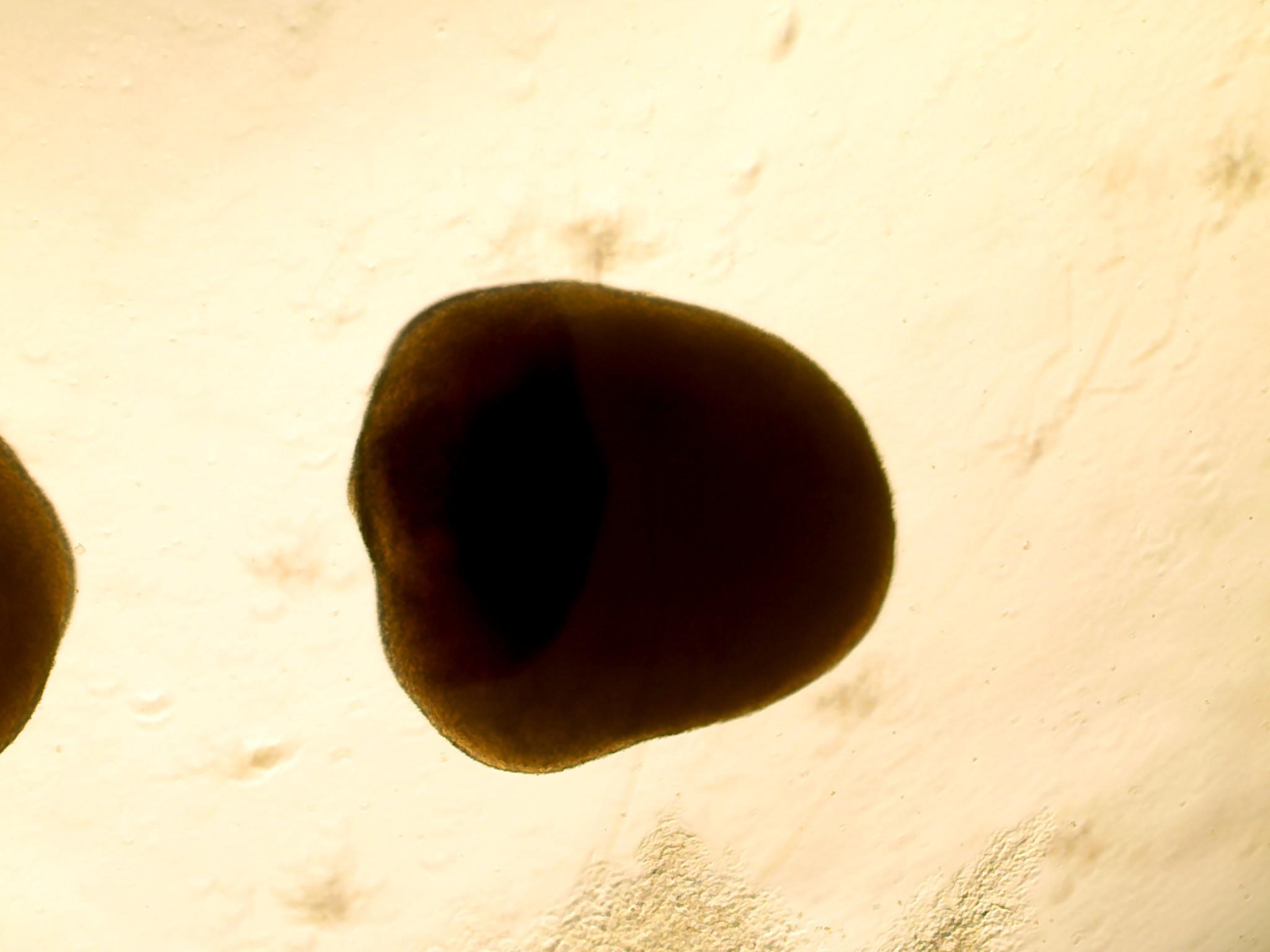 |
| --- | --- |
| hCO with unexpected morphology | 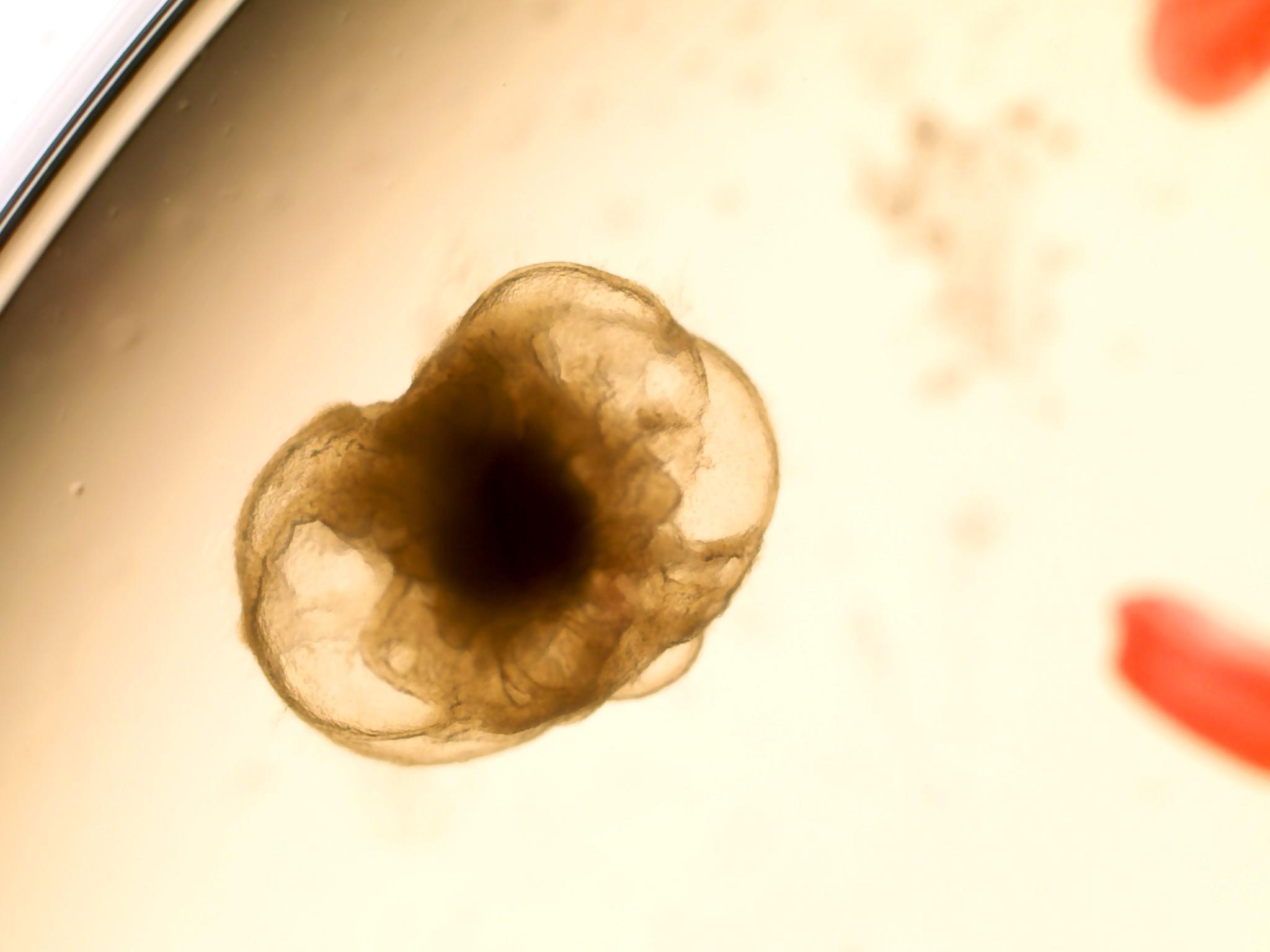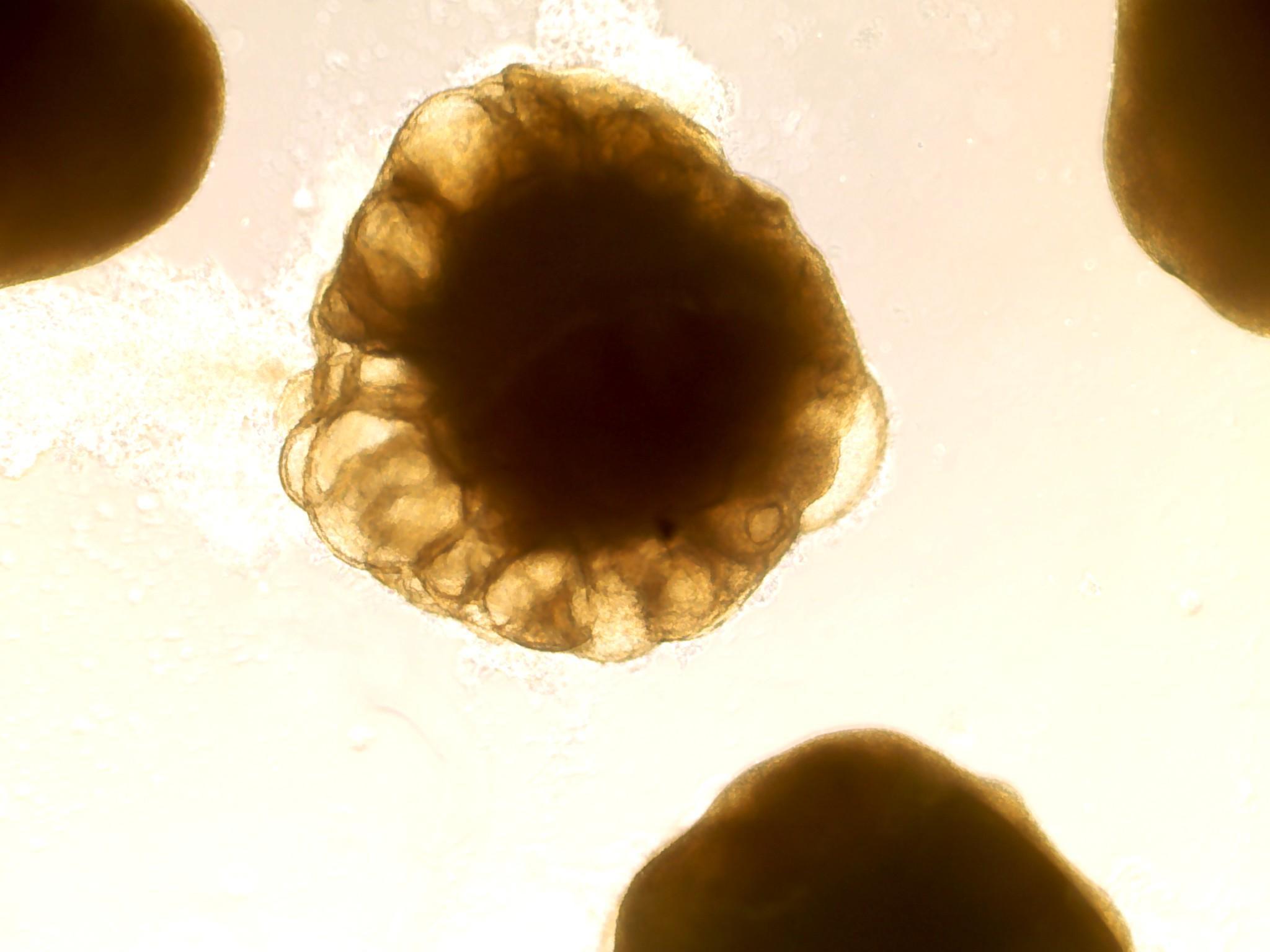 |

**References**:

Alves, Chrystian J., Rafael Dariolli, Frederico M. Jorge, Matheus R. Monteiro, Jessica R. Maximino, Roberto S. Martins, Bryan E. Strauss, José E. Krieger, Dagoberto Callegaro, and Gerson Chadi. 2015. “Gene Expression Profiling for Human iPS-Derived Motor Neurons from Sporadic ALS Patients Reveals a Strong Association between Mitochondrial Functions and Neurodegeneration.” *Frontiers in Cellular Neuroscience* 9 (August): 289.

Collier, Amanda J., Sarita P. Panula, John Paul Schell, Peter Chovanec, Alvaro Plaza Reyes, Sophie Petropoulos, Anne E. Corcoran, et al. 2017. “Comprehensive Cell Surface Protein Profiling Identifies Specific Markers of Human Naive and Primed Pluripotent States.” *Cell Stem Cell* 20 (6): 874–90.e7.

Watanabe, Momoko, Jessie E. Buth, Jillian R. Haney, Neda Vishlaghi, Felix Turcios, Lubayna S. Elahi, Wen Gu, et al. 2022. “TGFβ Superfamily Signaling Regulates the State of Human Stem Cell Pluripotency and Capacity to Create Well-Structured Telencephalic Organoids.” *Stem Cell Reports* 0 (0). https://doi.org/[10.1016/j.stemcr.2022.08.013](http://dx.doi.org/10.1016/j.stemcr.2022.08.013).
